# Exclusion Systems Preserve Host Cell Homeostasis and fitness, Ensuring Successful Dissemination of Conjugative Plasmids and Associated Resistance Genes

**DOI:** 10.1101/2025.04.18.649494

**Authors:** Agathe Couturier, Nathan Fraikin, Christian Lesterlin

## Abstract

Plasmid conjugation is a major driver of antibiotic resistance dissemination in bacteria. In addition to genes required for transfer and maintenance, conjugative plasmids encode exclusion systems that prevent host cells from acquiring identical or redundant plasmids. Despite their ubiquity, the biological impact of these systems remains poorly understood. Here, we investigate the importance of the exclusion mechanism for plasmid dynamics and bacterial physiology at the single-cell level. Using real-time microscopy, we directly visualize how the absence of exclusion results in plasmid unregulated self-transfer, causing continuous and repeated plasmid exchange among host cells. This runaway conjugation severely compromises cell integrity, viability, and fitness, a largely undescribed phenomenon termed lethal zygosis. We demonstrate that lethal zygosis is associated with membrane stress, activation of the SOS response and potential reactivation of SOS-inducible prophages, as well as chromosome replication and segregation defects. This study highlights how exclusion systems maintain host cell homeostasis by limiting plasmid transfer. Paradoxically, this restriction is critical to the successful dissemination of conjugative plasmids by conferring a selective advantage, which explains their evolutionary conservation and underscores their role in the spread of antibiotic resistance among pathogenic bacteria.

## INTRODUCTION

Conjugation is a horizontal gene transfer mechanism through which recipient bacteria acquire genetic information by direct contact from donor bacteria. Initially described with the discovery of genetic exchange in bacteria, which involved the F (fertility) factor, an extrachromosomal DNA element capable of autonomous replication and transfer between *Escherichia coli* cells (Lederberg and Tatum, 1946). Since then, plasmid conjugation has been revealed as a widespread mechanism of horizontal gene transfer, playing a central role in promoting genetic diversity and adaptation within microbial populations (Bottery, 2022; Tokuda and Shintani, 2024). Conjugative plasmids are recognized as the main contributors to the global spread of antibiotic resistance among bacterial pathogens, raising significant concerns about their impact on public health (Botelho and Schulenburg, 2021; Davies and Davies, 2010; MacLean and San Millan, 2019; Partridge et al., 2018). In particular, conjugative plasmids belonging to the IncF incompatibility group are responsible for the worldwide dissemination of resistance and virulence traits among clinical isolates of *Escherichia coli* and other enterobacterial species (Stephens et al., 2020).

Conjugation in Gram-negative bacteria primarily relies on plasmid-encoded factors involved in the key steps of DNA transfer, including the relaxosome protein complex that prepares the single-stranded DNA (ssDNA) plasmid before transfer, and the Type IV Secretion System (T4SS) that transports the ssDNA plasmid through the bacterial membrane and the sex pili that mediates mating pair formation (Mpf) (Costa et al., 2023; Fraikin et al., 2024; Virolle et al., 2020). Another conserved feature of conjugative plasmids is exclusion systems, encoded by diverse conjugative elements, including IncF, IncP, IncN, IncW, IncI, and IncH plasmids, mobilizable plasmids, integrative and conjugative elements (ICEs), as well as conjugative elements found in Gram-positive bacteria (Achtman et al., 1977; Garcillán-Barcia and de la Cruz, 2008; Igler et al., 2021; Rivard et al., 2024). Exclusion systems prevent the redundant acquisition of conjugative plasmids into a host cell that already contains identical or a closely related element. They achieve this by making the plasmid-carrying cell a poor recipient for conjugation, thereby reducing its efficiency of plasmid uptake. Although the precise molecular mechanisms underlying exclusion remains elusive, exclusion is known to be mediated by plasmid-encoded membrane proteins, which are classified into two groups based on their mode of interference in the conjugation process. In the F-like plasmids, surface exclusion factors (the outer-membrane protein TraT) inhibit mating pair formation (Mpf), while entry exclusion factors (the inner-membrane protein TraS) block DNA translocation following Mpf (Audette et al., 2007; Jalajakumari et al., 1987; Manning et al., 1980; Minkley and Ippen-Ihler, 1977; Seddon et al., 2025).

Early observations underscore the importance of exclusion for plasmid stability, reporting that F plasmid mutants harboring *traS* and *traT* mutations can never isolated by chemical mutagenesis suggesting that self-transfer-proficient mutants become highly unstable (Achtman et al., 1980, 1977). Furthermore, early genetic investigations have revealed the role of exclusion in protecting cells from lethal zygosis, a phenomenon initially described during mating with a large excess of Hfr (High frequency of recombination) donor cells capable of transferring the entire 4,6 Mb chromosome, which severely compromises the viability of the recipient cells (L. Alfoldi et al., 1957; Skurray and Reeves, 1973). These findings prompted the proposal that exclusion systems serve to shield host cells from excessive conjugation, which can disrupt essential cellular processes. Two main mechanisms have been proposed to explain lethal zygosis. First, the physical attachment of the conjugative pilus to the cell surface, along with the establishment of a complex protein channel through the cellular envelopes during mating may affect the integrity of the cell membranes. Second, superinfection by multiple copies of the ssDNA plasmid or increased copy number of double-stranded (dsDNA) plasmids could deplete the recipient cell’s metabolic or DNA processing resources and overwhelm the host cell resources. Both mechanisms have received some support from early studies. The addition of the outer membrane-disrupting antibiotic polymyxin B to the mating mix reduced the viability of transconjugant cells (Viljanen, 1987), while the specific inhibition of DNA transfer using nalidixic acid was found to partially suppress lethal zygosis (Ou, 1980). Despite these early proposals, the precise mechanisms underlying the phenomenon of lethal zygosis are still uncharacterized, and the specific physiological significance of exclusion systems remains largely unknown. To address this gap in understanding, we investigate the role of exclusion systems on plasmid dynamics and host cell physiology at the single-cell level.

## RESULTS

### Inactivation of exclusion triggers unrepressed plasmid self-transfer between host cells

The efficiency of exclusion is quantitatively assessed by estimating the exclusion index (EI), calculated by comparing the frequency of plasmid acquisition by a recipient strain harboring the plasmid to the frequency observed in the isogenic plasmid-free strain (Achtman et al., 1977). The wild-type F (F*wt*) plasmid, which carries the exclusion genes *traS* and *traT*, exhibits an EI of ∼474. This means that *E. coli* cells containing the F*wt* plasmid are 474 times less receptive to acquiring an additional plasmid than plasmid-free cells. Consistent with previous estimates, we calculate that single-deletion mutants lacking either *traT* (F*ΔtraT*) or *traS* (FΔ*traS*) exhibit reduced EIs of 28 and 3, respectively. Most importantly, we report for the first time the double deletion mutant (FΔ*traST*), which completely abolishes exclusion, as reflected by an EI of ∼1 (Fig. S1A). Exclusion is restored by complementation plasmids, both in the presence (Fig. S1B) and absence (Fig. S1C) of F plasmids, with TraS playing a more significant role than TraT. However, full exclusion requires the synergistic action of both proteins.

To investigate the impact of exclusion loss on plasmid dynamics and host cell physiology, we examined cells harboring the FΔ*traST* plasmid. In this clonal population, each cell is expected to function as both a plasmid donor and recipient. However, because all cells are genetically identical, traditional approaches based on resistance marker selection to measure plasmid transfer are not applicable. Therefore, we used live-cell microscopy coupled with established fluorescent reporters to visualize ssDNA plasmid transfer at the single-cell level (Couturier et al., 2023; Nolivos et al., 2019). Specifically, we used a fluorescent fusion of the chromosomally encoded single-strand binding protein Ssb-Ypet, and observed the frequent formation of bright membrane-associated foci on the ssDNA plasmid on both sides of the conjugation pore reflecting transfer between FΔ*traST*-carrying cells (Figure 1A; Movie S1). We estimate a dramatic increase in ssDNA plasmid frequency from 0.0063 transfers per cell per hour for F*wt* compared to 1.31 for FΔ*traST* (Figure 1B). This visualization approach provides the first direct observation of unrepressed ssDNA plasmid self-transfer among host cells triggered by the absence of exclusion system.

**Figure 1.**
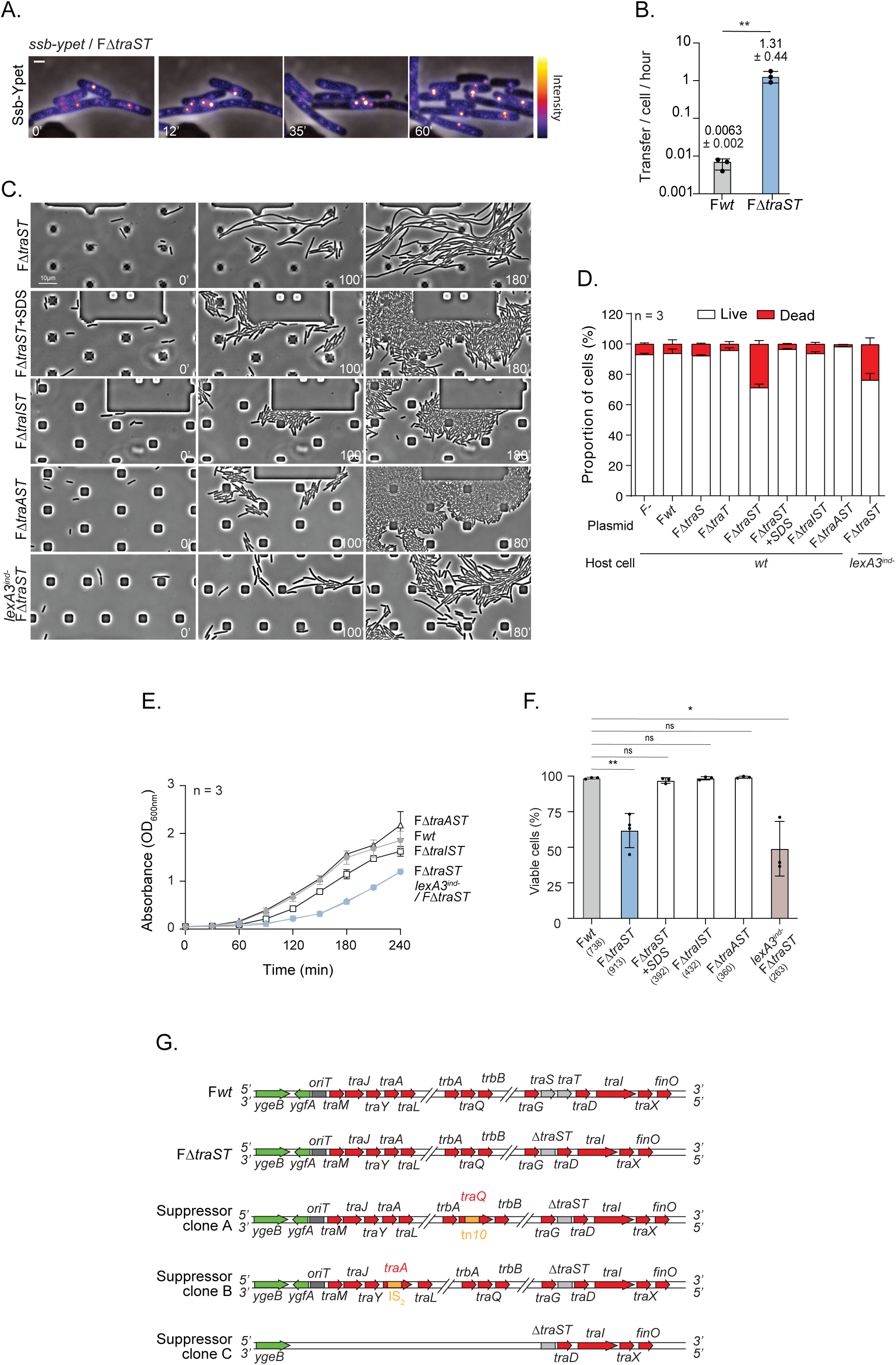
Loss of exclusion leads to deregulated plasmid transfer and impaired host cell viability. **(A)** Time-lapse microscopy images of a clonal population of FΔ*traST* cells carrying the *ssb-ypet* fusion gene, illustrating the transfer of single-stranded DNA (ssDNA). The intensity of Ssb-Ypet foci is represented by the color scale bar on the right. Scale bar: 1 µm. **(B)** Histogram showing the number of plasmid transfer events per cell per hour for cells carrying either F*wt* or F*ΔtraST*. Data represent mean and standard deviation (SD) from (n) individual transfer events across at least three independent biological replicates (black dots). Statistical significance was assessed using an unpaired t-test (\*\**P* = 0.0072). **(C)** Time-lapse phase-contrast images showing the filamentation phenotype in clonal populations of F*ΔtraST,* F*ΔtraST* cells treated with SDS, F*ΔtraIST,* F*ΔtraAST* and, *lexA3^ind-^* F*ΔtraST*. Scale bar: 10 µm. **(D)** Histograms showing the proportion of live and dead in F*wt,* F*ΔtraS,* F*ΔtraT* strains and its F*ΔtraST* plasmid derivatives, as determined by the Live/Dead staining assay. Bars represent mean and standard deviation (SD) from at least three independent experiments. **(E)** Growth curves of F*wt* and FΔ*traST* mutant cells, measured by OD600 every 30 minutes over 240-minutes. Cells were grown in LB medium in triplicate, only the average curve is shown. **(F)** Histograms showing the percentage of viable cells, defined by their ability to divide during time-lapse microscopy. Bars represent the mean ± SD from at least three independent biological replicates. **(G)** Schematic representation of the F*wt* and FΔ*traST* transfer regions and the corresponding genetic maps of the three suppressor clones (A, B and C).

### Unrepressed plasmid self-transfer induces host cell viability defects

Monitoring bacterial growth within a microfluidic chamber over 180 minutes using phase-contrast imaging shows that FΔ*traST-*carrying cells exhibit severe morphology defects, including the formation of filamentous and ghost cells (Figure 1C; Movie S2). Using live/dead staining and microscopy snapshot analysis, we quantified approximately 29% dead cells in the population (Figure 1D). These phenotypes result in growth defects and decreased viability, as demonstrated by OD monitoring (Figure 1E) and microscopy-based microcolony assays (Figure 1F), respectively. These deficiencies are observed in complete absence of exclusion genes, while only minor or no effects are observed in the *traS* or *traT* single mutants (Figure 1D and Fig.S1D). We further show that F*ΔtraST-*carrying cells exhibit increased uptake of SYTOX staining dye (FigS1E), which penetrates cells with compromised plasma membranes, and DiBAC_4_^(3)^ staining dye, which penetrate depolarized cells with impaired transmembrane potential (FigS1D). Such alterations of membrane permeability are likely related to the formation of ghost cells (Figure 1C-1D).

Crucially, cell filamentation and death, viability and growth defects induced by the absence of exclusion systems are suppressed when conjugation is abolished, either chemically by adding SDS to the microfluidic chamber, or genetically by deleting the relaxase gene *traI* or the pilin gene *traA*, which are essential for plasmid processing and transfer, and pilus formation, respectively (Figure 1C-F; Movie S2). This indicates that derepressed plasmid self-transfer is the sole cause of the observed proliferation defects. Consistent with this interpretation, suppressor mutants of the FΔ*traST*-carrying strain that regained normal growth in liquid conditions (Fig. S1G) have acquired mutations that drastically reduced or eliminated their capacity to transfer the FΔ*traST* plasmid (Fig. S1H). These mutations are found in the *tra* region, including a Tn*10* insertion interrupting the *traA* pilin gene essential for pilus formation, an IS2 insertion disrupting the *traQ* gene involved in pilin maturation, and a 22.85 kb deletion encompassing the origin of transfer (*oriT*) and part of the *tra* operon (Figure 1G). Altogether, these results demonstrate that unrepressed plasmid self-transfer results in severe impairment of cell integrity and viability, which imposes an important fitness burden to host cells.

### Unrepressed plasmid self-transfer induces membrane stress

In search for the mechanism by which unrepressed self-transfer impacts the physiology of the cells, we addressed its effect on the induction of five envelope stress response (ESR) pathways, namely Cpx, Rcs, Psp, SigmaE, and Bae. These ESR systems are known to be activated by specific perturbations of the cell envelope (Delhaye et al., 2019; Rousseau et al., 2023), i.e., the Cpx system by the presence of misfolded periplasmic proteins and by surface adhesion (Hunke et al., 2012; Raivio and Silhavy, 1997), Rcs by disturbances in the outer membrane (Majdalani et al., 2005), Bae by toxic compounds (Baranova and Nikaido, 2002; Raffa and Raivio, 2002), the phage-shock response Psp system by alterations in proton motive force (Brissette et al., 1990; Bury-Moné et al., 2009; Kleerebezem et al., 1996), while the envelope heat-shock sigma factor σ^E^ responds to the accumulation of unfolded outer membrane proteins or lipopolysaccharide alterations (Mitchell and Silhavy, 2019). We used transcriptional fusions coupling the promoters of genes specifically regulated by these ESRs (*cpxP*, *rcsA*, *pspA*, *micA*, and *mdtA*) to the gene encoding the fluorescent protein mNeonGreen (mNG) (Rousseau et al., 2023). The activation of these reporters was monitored in FΔ*traST*-carrying cell populations during exponential growth using single-cell microscopy analysis. No induction of Psp, Bae, or SigmaE pathways is observed in cells carrying F*ΔtraST* cells compared to cells carrying a F*wt* plasmid (Fig. S2A-C). However, we measured a significant induction of the Rcs (Figure 2A) and the Cpx (Figure 2B) pathways in F*ΔtraST-*carrying cells, indicative of a response to outer membrane perturbations and surface adhesion during unregulated self-transfer. Rcs and Cpx activations are mainly due to the absence of the surface exclusion protein *traT*, while weaker induction was observed in cells lacking entry exclusion (F*ΔtraS*) (Figure 2A). This suggests that stress pathways activation is related to processes occurring during the formation of the mating pair between cells. Consistently, Rcs and Cpx induction is fully abolished in F*ΔtraAST-*carrying cells, which are unable to form mating pair due to the lack of pili. However, Rcs and Cpx induction is also abolished by the deletion of the relaxase (F*ΔtraIST)* that retains pilus formation while impeding the processing of the plasmid required for transfer (Figure 2A-B). This suggests the activation of membrane stress pathways is due to interactions occurring between mating cells during repeated self-transfer of ssDNA plasmid through the conjugation pore. We measured a lesser but detectable induction of the Rcs pathway when conjugation is performed between F*wt* donor cells and *wt* recipients carrying the Rcs reporter (Fig. S2D). This indicates that outer membrane stress also occurs during normal conjugation, but is dramatically amplified by unrepressed self-transfer. Finally, we questioned whether Rcs and Cpx response pathways play any role in the viability of host cells acquiring plasmids DNA. However, we observed no effect of the deletion of these pathways on the proportion of dead cells in the presence or absence of exclusion (Figure 2C). Altogether, these results show that, while membrane stress are induced during conjugation, their downstream regulatory effect and biological functions are not critical to the survival of the host cells.

**Figure 2.**
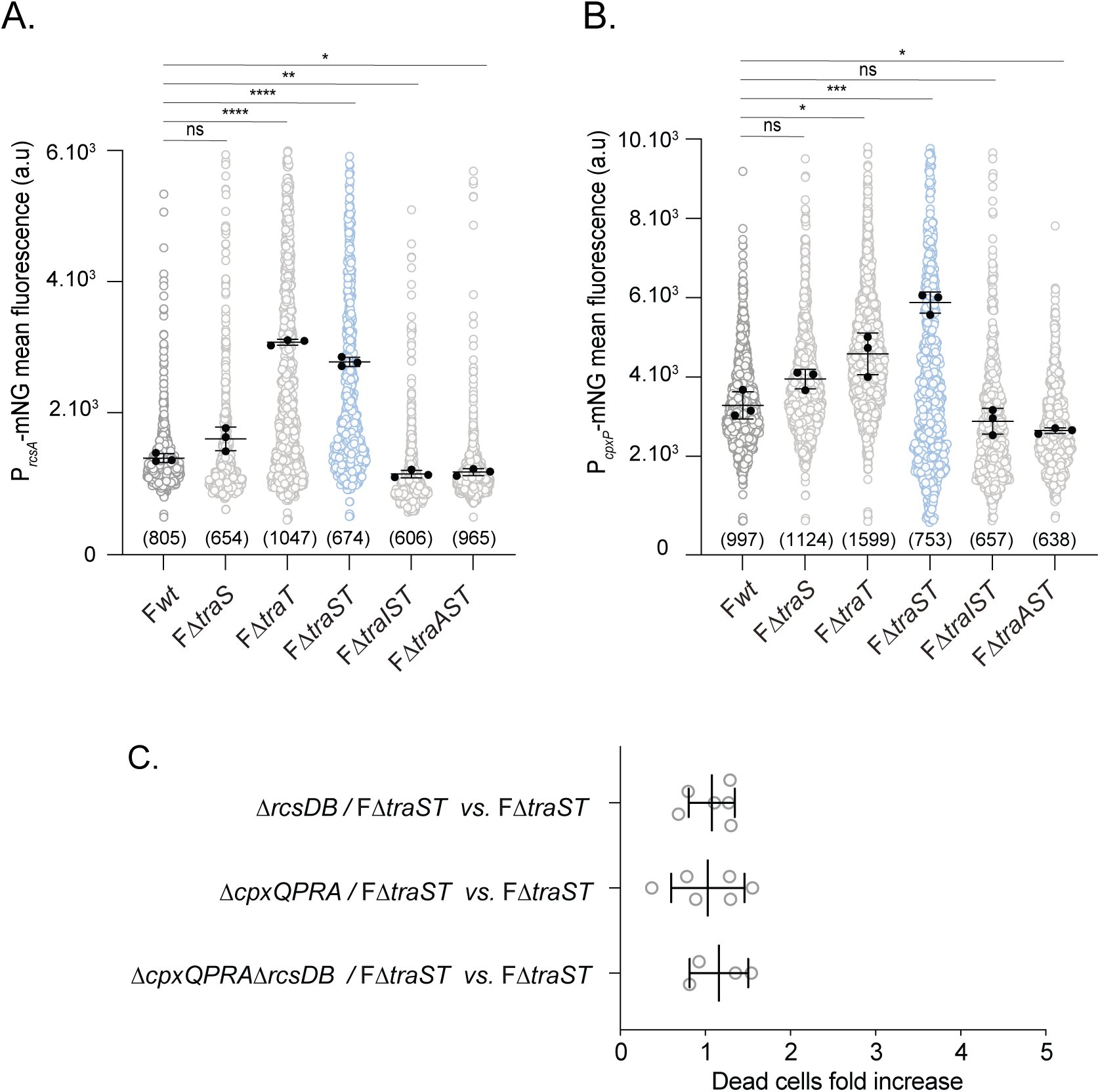
Exclusion systems limit envelope stress responses during plasmid transfer. **(A)** Quantification of P*_rcsA_*-mNG induction at the single-cell level in F*wt,* F*ΔtraS,* F*ΔtraT,* F*ΔtraST* and its derivative mutants during exponential growth. Each dot represents an individual cell; black dots indicate the mean fluorescence value for each independent biological replicate. Statistical significance was assessed using an unpaired t-test on replicate means (\**P* < 0.05*; **P* < 0.005*; ****P* < 0.0001). **(B)** Quantification of P*_cpxP_*-mNG induction at the single-cell level in F*wt,* F*ΔtraS,* F*ΔtraT,* F*ΔtraST* and its derivative mutants during exponential growth. Each dot represents an individual cell; black dots indicate the mean fluorescence value for each independent biological replicate. Statistical significance was assessed using an unpaired t-test on replicate means (\**P* < 0.05*; ***P* < 0.001). **(C)** Histograms showing the proportion of live and dead cells in strains carrying deletions in the Rcs (*ΔrcsDB*) *and/or* Cpx (*ΔcpxQPRA*) pathways, either without a plasmid (F-) or harboring F*ΔtraST.* Cell viability was assessed using the Live/Dead staining assay. Bars represent the mean ± standard deviation (SD) from at least three independent biological replicates.

### Unrepressed plasmid self-transfer induces activation of the SOS response

Next, we addressed whether the excessive entry of ssDNA during unregulated self-transfer induces the SOS response, possibly accounting for the observed viability and cell morphology defects. Using a transcriptional *sulA* fluorescent reporter (P*_sulA_*GFP), we measure a significant induction of the SOS response in F*ΔtraST*-carrying cells compared to cells containing F*wt* or no plasmid (Figure 3A). SOS induction is strictly attributable to unrepressed self-transfer, as it is fully suppressed when conjugation is impeded by the addition of SDS (Figure 3A) or by deletion of *traI* or *traA* (Fig. S3A). Consistent with this interpretation, we confirmed that the activation of the SOS in not due to the presence of DNA damage in F*ΔtraST*-carrying cells. To show this, we visualized a fluorescent fusion of the recombination protein RecA (RecA-YFP), which is diffuse in normally growing cells and form intracellular structures in response to DNA damage (Lesterlin et al., 2014; Wiktor et al., 2021). The formation of intracellular structures by fluorescent proteins can be reported by an increase in fluorescence skewness (Couturier et al., 2023; Shen et al., 2023). Analysis showed that while RecA-YFP skewness is significantly increased in response to UV-induced DNA damage (0.025 J/m²), it remained unchanged in F*ΔtraST* cells compared to F- and F*wt* cells (Fig. S3B).

**Figure 3.**
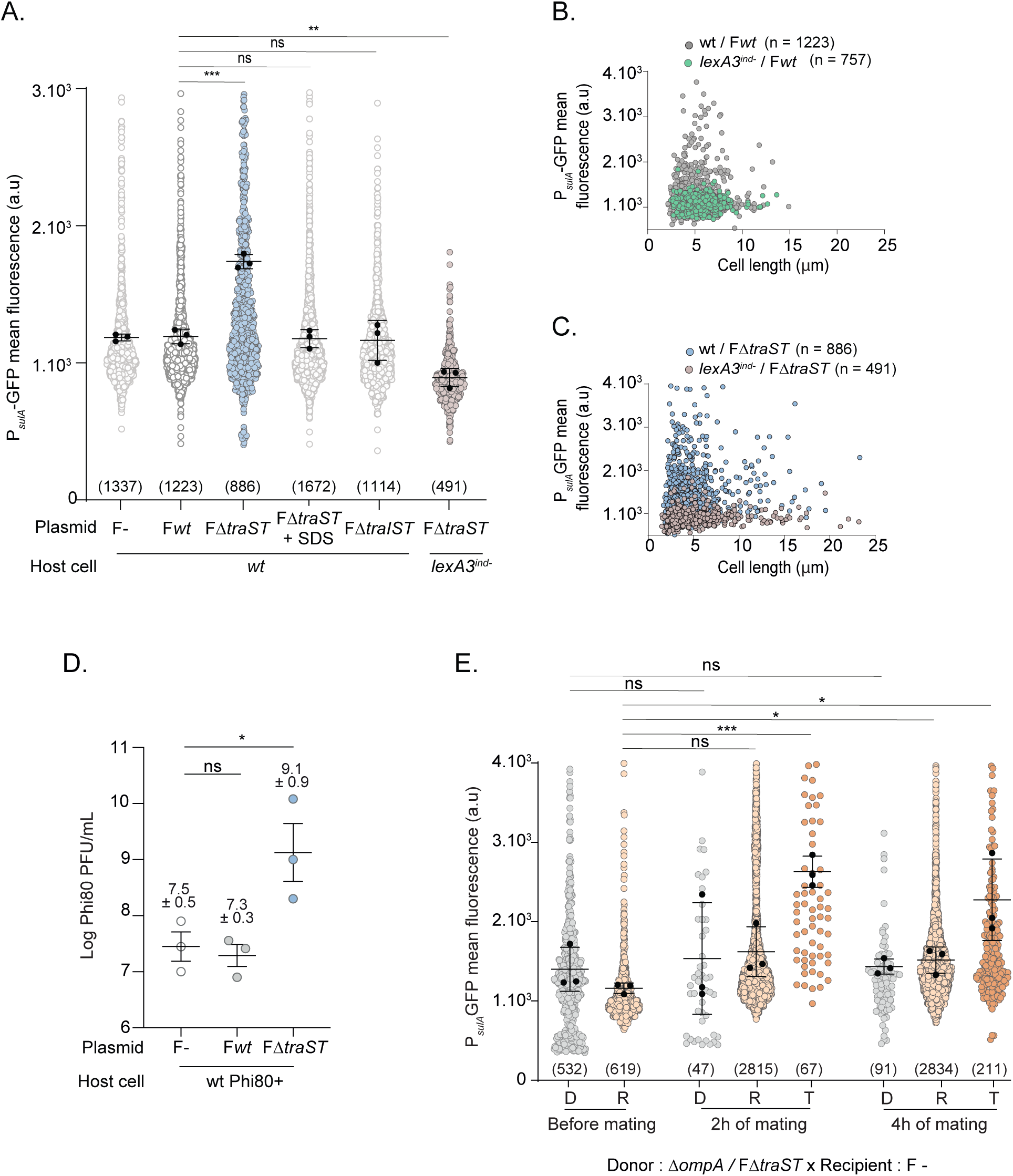
Induction of the SOS response and phage production upon exclusion system loss. **(A)** Quantification of P*_sulA_*-GFP reporter induction at the single cell level in F-, F*wt*, F*ΔtraST* or its derivatives. Each dot represents an individual cell; black dots indicate the mean fluorescence value for each independent biological replicate. Statistical significance was assessed using an unpaired t-test on replicate means (*\*\*P* < 0.005*; ***P* < 0.001). **(B)** Analysis of P*_sulA_*-GFP induction as a function of cell length in *wt* (grey circles) and *lexA3ind-* (green circles) strains carrying the F*wt* plasmid. The number of cells analyzed (n) is indicated, based on at least three independent experiments. **(C)** Analysis of P*_sulA_*-GFP induction as a function of cell length in *wt* (blue circles) and *lexA3ind-* (pink circles) strains carrying the F*ΔtraST* plasmid. The number of cells analyzed (n) is indicated, based on at least three independent experiments. **(D)** Quantification of Φ80 prophage production in F*wt,* F*ΔtraST* and plasmid free cells (F-). Data represent the mean ± SD from three independent experiments (black dots). Statistical significance was determined using a two-sided Mann–Whitney test (\**P = 0.0443; ns, not significant*). **(E)** P*_sulA_*-GFP intensity measured at the single cell level before mating, and after 2 and 4 hours of conjugation between a donor strain lacking *ompA* and carrying F*ΔtraST* plasmid and a plasmid-free recipient cell. Each dot represents an individual cell; black dots indicate the mean fluorescence value for each independent biological replicate.

To test the possibility that SOS activation could account for viability defects observed in the absence of exclusion, we examined the effect of the SOS-defective *lexA3^ind-^*mutation. While the *lexA3^ind^*mutation abolishes the activation of the SOS response in the F*ΔtraST* cells (Figure 3A), it does not suppress filamentation (Figure 1C, Figure 3B and 3C) or cell death (Figure 1D), and does not improve viability nor cell growth (Figure 1E and 1F). We therefore conclude that the onset of the SOS response is not responsible for the viability defects triggered by the absence of exclusion. However, we further hypothesized that the induction of strong levels of SOS response due to the loss of exclusion could result in the awakening of SOS-inducible prophage potentially located into the host chromosome. To test this possibility, we measured the concentration of phage in the supernatant of culture of *wt E. coli* strains carrying the Φ80 prophage, with or without F derivatives. We observed a 2-log increase in phage plaque-forming units (PFU/ml) in cultures with the F*ΔtraST* plasmid compared to those carrying F*wt* or F-cells (Figure 3D).

Our experimental setup gave us the unique opportunity to address whether, in the F*ΔtraST* population where each cell can act as both a donor and a recipient, the induction of SOS occurs within the donor due to plasmid donation or within the recipient due to plasmid acquisition, or both. To address that specific question, we used a *ΔompA* donor strain that has reduced capacity to receive the plasmid, as OmpA protein is critical to the mating pair formation (Manoil and Rosenbusch, 1982; Ried and Henning, 1987). The *ΔompA /* F*ΔtraST* donor retains plasmid donation proficiency (Fig. S3C), but has approximately 200-fold reduced plasmid acquisition capability (Fig. S3D). During conjugation, SOS induction is not observed in the *ΔompA /* F*ΔtraST* donor, but only in the subpopulation of transconjugant cells that have received the F*ΔtraST* plasmid (Figure 3E).

All together, these findings demonstrate that the activation of the SOS response in the absence of exclusion systems is due to repeated plasmid acquisition rather than repeated plasmid donation. However, SOS induction is not responsible for the viability defects induced by unrepressed plasmid self-transfer.

### Impact of the loss of exclusion on replicon maintenance

Next, we questioned whether the loss of exclusion could result in a deregulation of plasmid copies per cell or maintenance, which could in turn have deleterious impact on the physiology of the host cell. Indeed, an excess in plasmid copies per cell might overwhelms the cell’s resources and induce a dramatic fitness cost (Rouches et al., 2022). Hence, we addressed if the ssDNA plasmids acquired through unrepressed self-transfer can be successfully converted into double-stranded DNA (dsDNA) plasmids. To this end, we examined mating between two parental strains, one harboring the FΔ*traST* plasmid labeled with a green fluorescent sfGFP-ParB_P1_/*parS*_P1_ system (F^GFP+^ cells) and the other carrying a red fluorescent mCh-ParB_PMT1_/*parS*_PMT1_ system (F^mCh+^ cells). Upon mating between F*ΔtraST*^GFP+^ and F*ΔtraST*^mCh+^ cells, we observed the rapid emergence of dual-labeled cells containing both plasmids (Figure 4A; Movie S3), constituting ∼70% of the population after four hours of mating (Figure 4B). Notably, dual-labeled cells were rarely detected when the conjugation mix was treated with SDS, which impedes conjugation by depolymerizing the conjugative pilus (Figure 4B), or when the experiment was performed with F*wt* plasmids (Figure 4C). These observations indicate that ssDNA F*ΔtraST* plasmids that are transmitted among and between F*ΔtraST*-bearing cells can, at least partially, be successfully converted into dsDNA.

**Figure 4.**
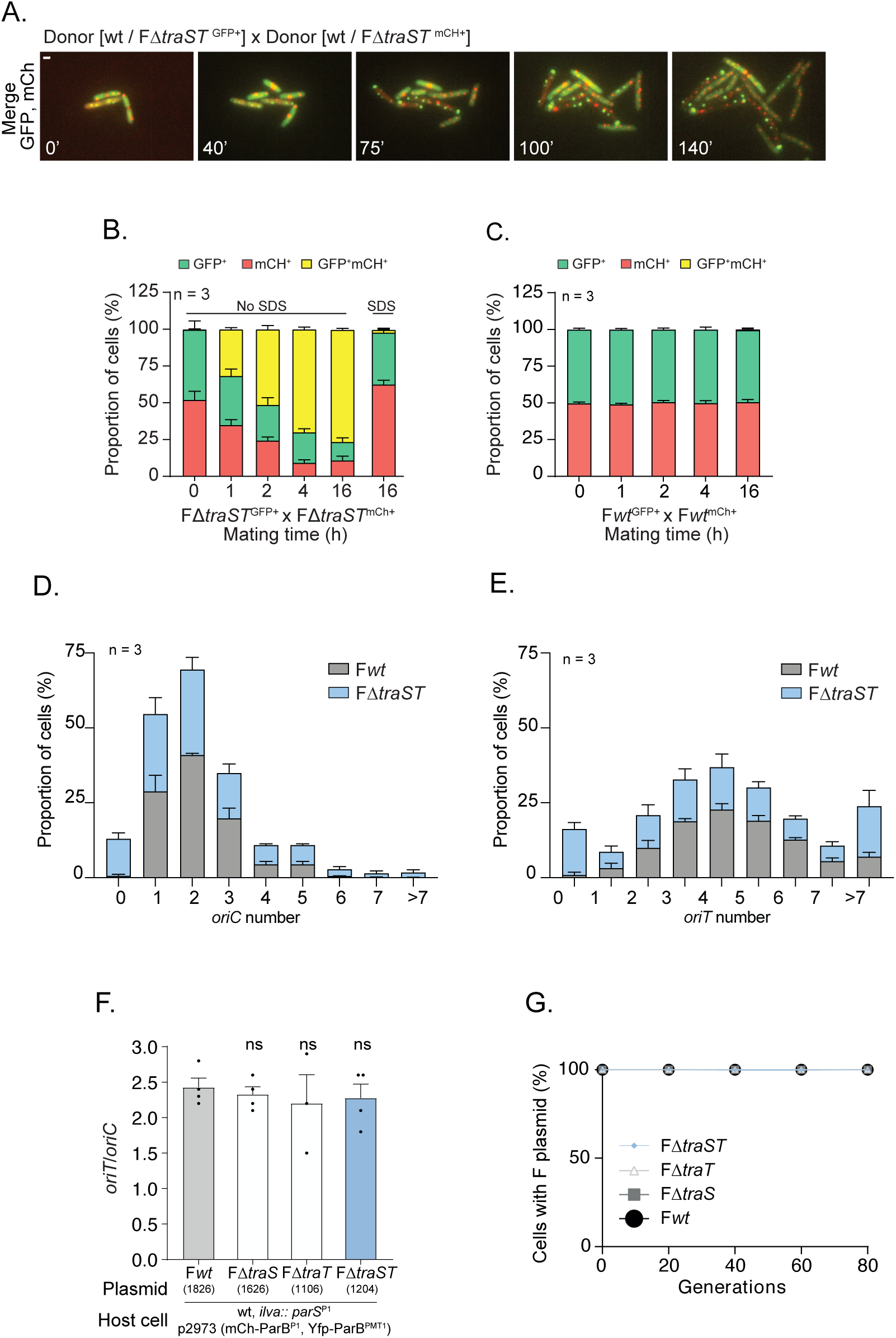
Impact of the loss of exclusion on replicon maintenance. **(A)** Time-lapse microscopy images showing the transfer of dsDNA in FΔ*traST*-carrying cells, indicated by the appearance of a GFP focus in an mCH+ cell or vice versa. Scale bar: 1 µm. **(B)** Quantification of plasmid transfer over time in mating assays between FΔ*traST^GFP+^* and FΔ*traST^mCH+^* strains. Bars represent the proportion of GFP^+^, mCH^+^ and double-positive GFP^+^mCH^+^ cells at the indicated time points, with or without SDS treatment. Data represent the mean and SD from three independent biological replicates (n=3). **(C)** Quantification of plasmid transfer over time in mating assays between F*wt^GFP+^* and F*wt^mCH+^* strains. Bars represent the proportion of GFP^+^, mCH^+^ and double-positive GFP^+^mCH^+^ cells at the indicated time points. Data represent the mean and SD from three independent biological replicates (n=3). **(D–E)** Histograms showing the distribution of *oriC* **(D)** and *oriT* **(E)** foci number per cell in populations carrying either the F*wt* or FΔ*traST* plasmid, as determined by fluorescence microscopy. **(F)** Histogram showing the *oriT*/*oriC* ratio in cells carrying either the F*wt*, F*ΔtraS,* F*ΔtraT* or F*ΔtraST* plasmid. The experiment was performed from at least three independent biological replicates. *P*-value significance ns were obtained from Mann–Whitney two-sided statistical test. **(E)** Stability of F*wt*, F*ΔtraS,* F*ΔtraT* or F*ΔtraST* plasmid under nonselective conditions. Plasmid retention rate was obtained from 20, 40, 60 and 80 generations of growth.

Because newly acquired ssDNA plasmids can be converted into dsDNA plasmids, we asked whether repeated self-transfer alters the intracellular copy number of the F plasmid. To test this, we quantified the *oriT*/*oriC* ratio in a strain carrying a *parS_P1_* insertion in the *ilvA* chromosome locus located near the origin of replication (*oriC*), and harboring the F plasmid with a *parS_PMT1_* insertion adjacent to the origin of transfer (*oriT*) (Fig. S4A and S4B). Quantitative image analysis reveals that the F*ΔtraST* population exhibit ∼10% of cells without *oriC* and *oriT* foci, a phenotype absent from the F*wt* control (Figure 4D and Figure 4E). Size distribution analysis showed that these focus-negative cells are predominantly small, consistent with recently divided newborn cells, which are likely non-viable and leading to the eventual formation of ghost cells (Fig. S4C). Nonetheless, most FΔ*traST* cells are normal-length or filamentous and exhibit an *oriT*/*oriC* ratio of ∼2.4, similar from F*wt* cells, indicating that the chromosome-to-plasmid ratio is maintained in absence of exclusion (Figure 4F). Consistent with plasmid maintenance, stability assay shows that, similar to F*wt*, all viable cells retain FΔ*traST* plasmid when grown over 80 generations without selection pressure (Figure 4G).

### Unrepressed plasmid self-transfer induces cell cycle disruption

Next, we characterized the impact of superinfection with ssDNA plasmid on the host’s cell chromosome dynamics. In particular, we characterized chromosome intracellular positioning using DAPI staining, and replication intracellular organization using a fluorescent fusion of the replisome’s B2 subunit (mCh-DnaN). In F*wt*-containing cells, DAPI-fluorescence demographs of cells sorted by length and corresponding localization heatmaps reflect the progressive segregation of well-separated nucleoid DNA in the course of cell cycle (Figure 5A). By contrast, F*ΔtraST*-containing cells exhibit impaired nucleoid separation, especially marked in small cells (<4 µm) (Figure 5A) as well as in filamentous (> 8 µm) cells (Fig. S5A). This phenotype is associated with mislocalization of the replication machinery. In F*wt*-containing cells, mCh-DnaN forms discrete foci regularly positioned at the cell quarter positions, reflecting replisome localization during DNA replication progression (Figure 5B). By contrast, localization analysis of F*ΔtraST* cells revealed a dramatic impairment of mCherry-DnaN localization, which appear largely diffuse in smaller cells (< 4µm) and strongly mispositioned foci in larger cells (> 4µm) (Figure 5B), as well as in filamentous cells (Fig. S5B). In parallel, we quantified the cell DNA content using Syto9 staining and flow cytometry (Figure 5C). The results show that most F*ΔtraST*-carrying cells have significantly lower DNA content compared to those carrying the F*wt* plasmid, reflecting a significant disruption of chromosome replication.

**Figure 5.**
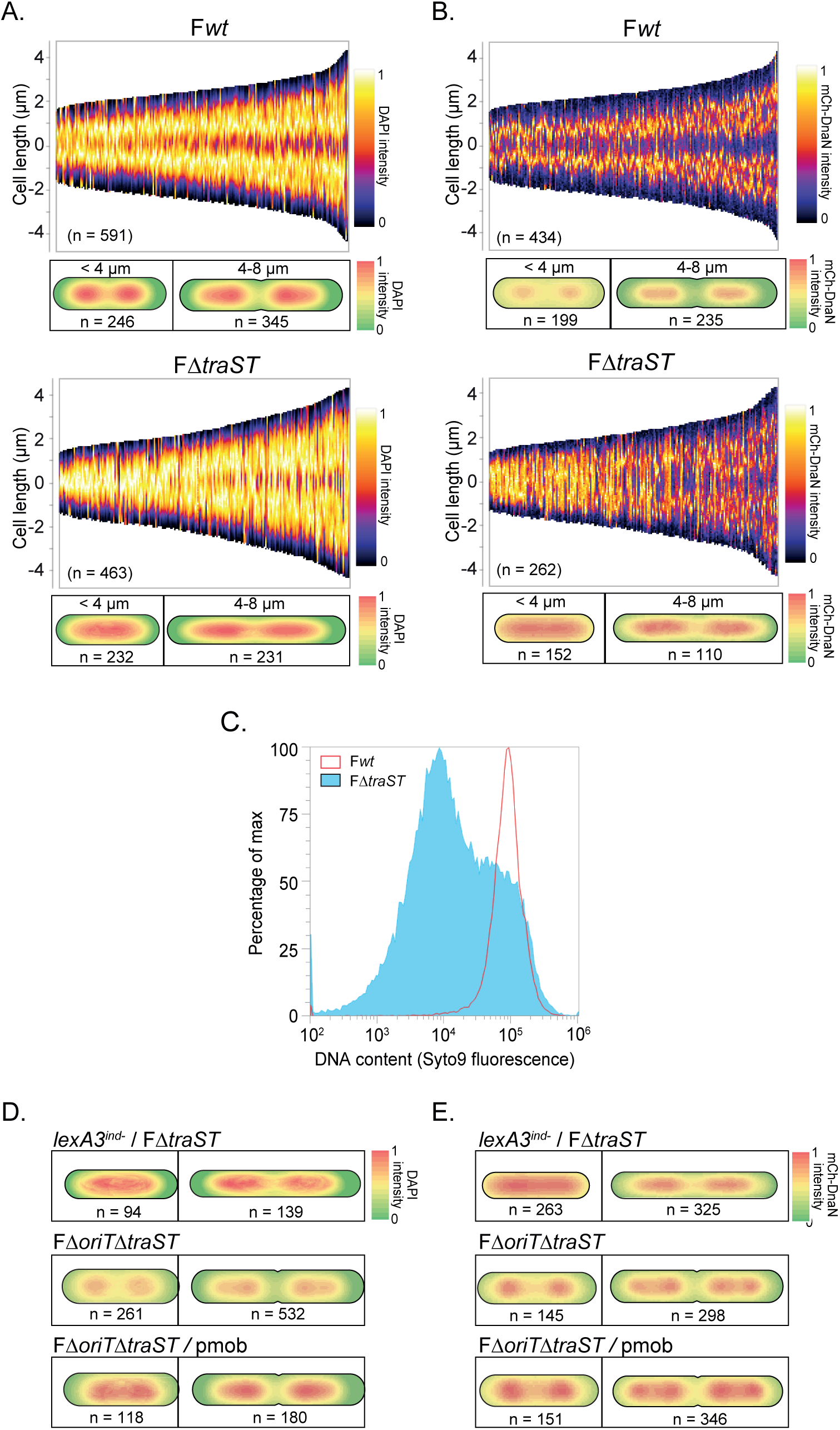
Effect of transferred DNA on chromosome organization and replication dynamics. **(A)** Nucleoid positioning visualized by DAPI staining in strains carrying F*wt* or F*ΔtraST*. **(B)** Replisome localization using a mCh-DnaN fusion in the same strains. For both panels, demographs show signal distribution in cells sorted by length (4–8 µm), and heatmaps display average fluorescence intensity along the normalized cell axis for short (<4 µm) and elongated (4–8 µm) cells. The number of analyzed cells (n) is indicated, based on data from three independent experiments. **(C)** Quantification of DNA content using Syto9 staining in cells carrying F*wt,* and F*ΔtraST.* **(D)** Heatmaps showing the spatial distribution of DAPI-stained nucleoids and replisome localization using mCh-DnaN fusion **(E)** in exponentially growing cells. Cells are grouped by size: small (<4 µm) and normal-sized (4–8 µm). Data are shown for clonal populations of *lexA3^ind-^ /* F*ΔtraST, wt /* FΔ*oriTST* and wt / FΔ*oriTST* / pmob. The number of analyzed cells (n) is indicated, based on data from three independent experiments.

Nucleoid and replisome mislocalizations persist in *lexA3^ind-^*cells (Figure 5C and 5D, and Fig. S5C and S5D), further supporting our previous interpretation that the SOS response does not account for the defects associated with the absence of exclusion. We then wanted to know if the amplitude of these defects depends on the amount of DNA transferred. To test this, we first abolished FΔ*traST* plasmid transfer by deleting its origin of transfer (FΔ*oriTtraST*), which restores normal chromosome and replisome dynamics (Figure 5C and 5D), suppresses SOS induction as measured using the *sulA* transcriptional fluorescent reporter (Fig. S5E), and prevents the formation of dead cells as assessed by live and dead staining (Fig. S5F). We then introduced a 6 kb plasmid (pmob) that carries the *oriT* from the F plasmid, rendering it mobilizable by the conjugation machinery produced from the co-resident FΔ*oriTtraST* plasmid. We observe that the reduction of the size of the transferred DNA from 108 kb of the F plasmid to the 6 kb of the pmob plasmid significantly reduces SOS induction and suppresses the formation of dead cells (Fig. S5E and S5F). Most importantly, it significantly improves nucleoid separation and restores normal replisome localization (Figure 5C and 5D). Conversely, we attempted to increase the amount of transferred DNA by removing exclusion systems from an Hfr strain that is able of transferring its entire 4,6 Mb chromosome. However, despite multiple attempts, we were unable to construct a viable deletion strain, further suggesting that unregulated self-transfer with extreme DNA amounts imposes a lethal burden on host cells. These findings are consistent with the interpretation that the amount of transferred DNA is a key determinant of the cellular stress induced by conjugation.

### Exclusion safeguard successful plasmid dissemination

The absence of exclusion systems and the consequent unrepressed self-transfer impacts negatively the fitness of the host cell. We wanted to evaluate how these fitness defects influence the dissemination capability of a plasmid devoid of exclusion system within a bacterial population. To address this question, we performed competition assays by mixing equal proportions of cells carrying FΔ*traST*^RFP+^ (*RFP* fusion) and cells carrying F*wt*^GFP+^ (*GFP* fusion). We then monitored their relative ratio of over several time scales. Using fluorescence microscopy, we observe that the proportion of cell carrying FΔ*traST* relative to F*wt* decreases from 1:1 to 1:13 over the first 20 hours (Figure 6A). In comparison, competition between two F*wt* populations showed no significant change in their relative proportions (Figure 6A). This trend was further confirmed after 24 hours by fluorescence imaging of bacterial colonies derived from the initial competition mix, revealing a clear reduction in the FΔ*traST*^RFP+^ signal relative to the F*wt*^GFP+^ (Figure 6B). To investigate this effect on longer period, we monitored the competition mix over a week using platting assays. F*wt* rapidly outcompetes FΔ*traST*, reaching up to a 5-log advantage after seven days (Figure 6C). These experiments demonstrate that exclusion systems are critical to the successful dissemination and maintenance of the plasmid within a bacterial population.

**Figure 6.**
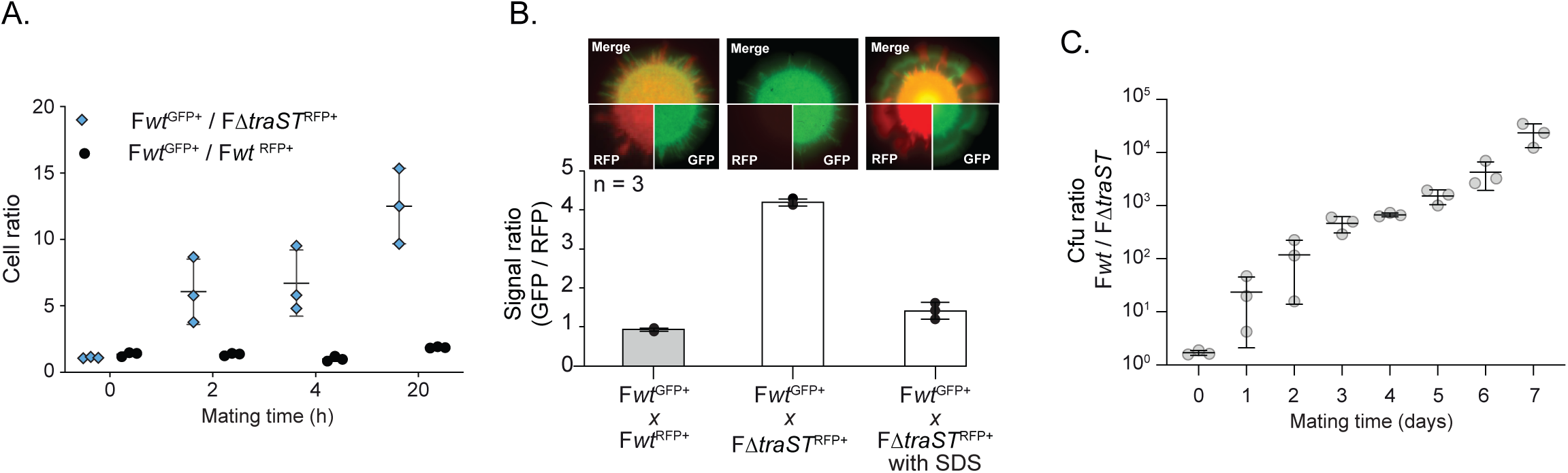
Exclusion safeguard successful plasmid dissemination. **(A)** Quantification of plasmid ratios within cell population during mating between F*wt*^GFP+^ and either F*ΔtraST*^RFP+^ or F*wt*^RFP+^. Ratios were estimated by fluorescence microscopy at 0, 2, 4 and 20 hours post-mating, across three independent experiments. **(B)** Quantification of the GFP/RFP fluorescence signal ratio after 24 hours of mating between F*wt*^GFP+^ and either F*ΔtraST*^RFP+^ or F*wt*^RFP+^, with and without SDS treatment. Representative merged fluorescence images are shown for each mating. Data represent the mean ± SD from three independent experiments (n = 3). **(C)** Quantification of plasmid ratios over time between F*wt* and F*ΔtraST* strains during co-culture. The ratio of colony forming units (CFU) was measured daily over a 7-day mating experiment.

## DISCUSSION

This study demonstrates that exclusion systems play a crucial role in maintaining a healthy host population capable of effectively propagating plasmids by mitigating the harmful effects associated with unregulated plasmid transfer and resulting in lethal zygosis. While this conclusion aligns with previous proposals (Davis and Grossman, 2021; Garcillán-Barcia and de la Cruz, 2008; Ou, 1980), our findings provide a detailed description of the events occurring during lethal zygosis. In the absence of functional exclusion mechanisms, host cells simultaneously act as efficient donors and recipients, initiating a runaway conjugation process characterized by frequent and continuous ssDNA plasmid exchanges. This unrestricted self-transfer profoundly impacts host physiology, causing filamentation, cell death, reduced viability, and significant fitness defects.

A direct consequence of excessive conjugation is the induction of membrane stress and activation of the Rcs and Cpx phosphorelay systems. Several steps within the conjugation process can compromise membrane integrity, including physical interactions mediated by the conjugative pilus, stabilization of mating pairs via OmpA–TraN interactions (Frankel et al., 2023; Low et al., 2023, 2022), channel formation through the cell envelope by the T4SS, and the passage of ssDNA through the conjugation pore. However, our observations that neither Rcs nor Cpx inactivation affected transconjugant cell viability during either regulated or runaway conjugation strongly suggest that these pathways do not significantly contribute to maintaining host cell homeostasis during plasmid acquisition. Furthermore, membrane stress responses depends primarily on the absence of the surface exclusion protein TraT, which interferes with OmpA-TraN interaction (Riede and Eschbach, 1986), and were entirely suppressed by traA deletion. Thus, membrane stress predominantly arises from events occurring during or after mating pair formation. Inactivation of the *traI* gene, which impedes plasmid transfer but does not affect pilus production, suppresses membrane stress activation and fully prevents filamentation, cell death, and growth defects. This finding supports the conclusion that the harmful effects on host physiology primarily result from to the repeated passage of the ssDNA plasmids through the membranes-spanning conjugation pore rather than by pilus-mediated interactions *per se*. We then investigated several potential consequences of superinfection by excessive ssDNA.

We report that unrepressed ssDNA entry triggers the activation of the LexA-dependent SOS response. Our data reveals that unrepressed ssDNA transfer strongly induces the SOS response, specifically due to repeated plasmid acquisition rather than donation. However, SOS induction itself does not explain the proliferation defects observed during unregulated self-transfer. Nonetheless, SOS activation led to secondary effects, notably the reactivation of SOS-inducible prophages. This can lead to further implications, as reactivated prophages can trigger host cell lysis and alter competition among mobile genetic elements, potentially favoring certain plasmids or mobile elements over others.

The absence of exclusion does not result in an increase in FΔ*traST* plasmid copy number per cell, eliminating resource overload as a cause of fitness defects. The maintenance of FΔ*traST* copy-number supports the model that, despite repeated ssDNA acquisition, the regulation of F-plasmid by host-chromosome replication preserves a constant plasmid-to-chromosome ratio (Keasling et al., 1992), further suggesting that not all incoming ssDNA molecules are converted into dsDNA plasmids The absence of exclusion also results in the formation of a significant proportion of cells that have lost at least part of their chromosome or plasmid DNA content. While that these cells are expected to degenerate into ghost cells, it is more difficult to determine their mechanism of formation. Two mechanisms could theoretically generate these DNA-deficient cells, i.e., the asymmetric polar division of filaments that results in cells with aberrant DNA content (Cayron et al., 2023; Raghunathan et al., 2020), or post-segregational killing (PSK) triggered by the activation of toxin–antitoxin modules upon plasmid loss (Fraikin and Van Melderen, 2024). However, multiple lines of evidence demonstrate that PSK does not account for the growth defects triggered by plasmid self-transfer. First, both normal and filamentous FΔ*traST* cells retain their plasmids, ruling outkilling through toxin–antitoxin activation after plasmid loss. Second, chemically blocking conjugation with SDS or genetically deleting *traA* or *traI* genes fully restores normal growth. Taken together, these observations implicate runaway conjugation itself as the cause of proliferation deficiencies.

We report that a major consequence of superinfection with ssDNA plasmids is the significant disruption of the intracellular organization of host chromosome replication and segregation. Based on these observations, we propose that the disruption of chromosome dynamics is primarily due to persistent recruitment of essential host factors such as single-stranded DNA-binding protein (Ssb), DNA polymerase III (DnaE) that to converts the ssDNA plasmids into double-stranded DNA (Wilkins and Hollom, 1974; Willetts and Wilkins, 1984) and potentially other replication machinery components onto incoming ssDNA plasmids. The hijacking of these essential factors would consequently limit their availability for chromosome maintenance, explaining the observed chromosome replication defects, reduced DNA content, and impaired cell proliferation. Similar stresses may also arise transiently during regulated conjugation, when functional exclusion systems limit ssDNA entry. Although brief and moderate, these effects likely contribute to the immediate plasmid-acquisition cost that reduces host fitness upon plasmid entry (Ahmad et al., 2023; Prensky et al., 2021). Under uncontrolled self-transfer, however, the same stresses accumulate continuously, eventually overwhelming cellular functions and imposing an insurmountable fitness burden.

Overall, this work sheds light on the pivotal role of exclusion systems in preserving host cell homeostasis and fitness. Although counterintuitive, restricting plasmid transfer actually enhances the plasmid’s dissemination potential by balancing the need for horizontal gene transfer against maintaining host viability. In evolutionary terms, plasmids that carry well-functioning exclusion systems gain a selective advantage, thereby promoting a more stable coexistence with their bacterial hosts and facilitating the dissemination of conjugative elements and associated antibiotic resistance genes among bacteria. These findings also emphasize the importance of further investigating the largely unknown molecular mechanisms underlying surface and entry exclusion systems. Targeting these systems could induce lethal zygosis in plasmid-bearing cells, opening a novel strategy to limit the spread of antibiotic resistance by conjugation.

## METHODS

### Bacterial strains, plasmids and growth

Bacterial strains are listed in Tables S1, plasmids in Table S2, and oligonucleotides in Tables S3. Fusion of genes with fluorescent tags and gene deletion on the F plasmid used λRed recombination (Yu et al., 2000; Datsenko and Wanner, 2000). Modified F plasmids were transferred to the background strain K12 MG1655 by conjugation. Where multiple genetic modifications on the F plasmid were required the kan and cat genes were removed using site-specific recombination induced by expression of the Flp recombinase from plasmid pCP20 (Datsenko and Wanner, 2000). Plasmid cloning were done by Gibson Assembly and verified by Sanger sequencing (Eurofins Genomics biotech). Strains and plasmids were verified by Sanger sequencing (Eurofins Genomics). Cells were grown at 37°C in M9 medium supplemented with glucose (0.2 %) and casamino acid (0.4 %) (M9-CASA) before imaging, and in Luria-Bertani (LB) broth for conjugation efficiency assays. When appropriate, supplements were used in the following concentrations; Ampicillin (Ap) 100 µg/ml, Chloramphenicol (Cm) 20 µg/ml, Kanamycin (Kn) 50 µg/ml, Streptomycin (St) 20 µg/ml, and Tetracycline (Tc) 10 µg/ml.

### Conjugation assays

Overnight cultures in LB of recipient and donor cells were diluted to an A600 of 0.05 and grown until an A600 comprised between 0.7 and 0.9 was reached. 25 µl of donor and 75 µl of recipient cultures were mixed into an Eppendorf tube and incubated for 90 minutes at 37°C. 1 ml of LB was added gently and the tubes were incubated again for 90 min at 37°C. Conjugation mix were vortexed, serial diluted, and plated on LB agar X-gal 40 µg/ml IPTG 20 µM supplemented the appropriate antibiotic to select for recipient or donor populations. Recipient (R) colonies were then streaked on plated on LB agar containing tetracycline 10 µg/ml to select for transconjugants (T) and the frequency of transconjugant calculated from the (T/R+T) presented in Figure 4C.

### Live-cell microscopy experiments

Overnight cultures in M9-CASA were diluted to an A600 of 0.05 and grown until A600 = 0.8 was reached. Conjugation samples were obtained by mixing 25 µl of donor and 75 µl of recipient into an Eppendorf tube. For time lapse experiments, 50 µl of the pure culture or conjugation mix was loaded into a B04A microfluidic chamber (ONIX, CellASIC®) (Cayron and Lesterlin, 2019). Nutrient supply was maintained at 1 psi and the temperature maintained at 37°C throughout the imaging process. Cells were imaged every 1 or 5 min for 90 to 120 minutes. For snapshot imaging, 10 µl samples of clonal culture or conjugation mix were spotted onto an M9-CASA 1% agarose pad on a slide (Lesterlin and Duabrry, 2016) and imaged directly.

Image acquisition. Conventional wide-field fluorescence microscopy imaging was carried out on an Eclipse Ti2-E microscope (Nikon), equipped with x100/1.45 oil Plan Apo Lambda phase objective, ORCA-Fusion digital CMOS camera (Hamamatsu), and using NIS software for image acquisition. Acquisitions were performed using 50% power of a Fluo LED Spectra X light source at 488 nm and 560 nm excitation wavelengths. Exposure settings were 100 ms for Ypet, sfGFP and mCherry and 50 ms for phase contrast.

Image analysis. Quantitative image analysis was done using Fiji software with MicrobeJ plugin (Ducret et al., 2016). For snapshot analysis, cells’ outline detection was performed automatically using MicrobeJ and verified using the Manual-editing interface. For time lapse experiments, detection of cells was done semi-automatedly using the Manual-editing interface, which allows to select for the cells to be monitored and automatically detect the cell outlines. Within conjugation populations, donor (no mCh-ParB signal), recipient (diffuse mCh-ParB signal), or transconjugant (mCh-ParB foci) category were assigned using the ‘Type’ option of MicrobeJ. Recipient cells were detected on the basis of the presence of red fluorescence above the cell’s autofluorescence background level detected in the donors. Among these recipient cells, transconjugants were identified by running MicrobeJ automated detection of the ParB fluorescence foci (Maxima detection). This approach was used independently of the presence or the absence of the Ssb-Ypet, or sfGFP fusions within donor and recipient cells. Within the different cell types, mean intensity fluorescence (a.u.), skewness, Signal/Noise Ratio (SNR), or cell length (µm) parameters were automatically extracted and plotted using MicrobeJ. SNR corresponds to the ratio (mean intracellular signal / mean noise signal), where the mean intracellular signal is the fluorescence signal per cell area and the noise is the signal measured outside the cells (due to the fluorescence emitted by the surrounding medium). By contrast with the total amount of fluorescence per cell, which is depending on the cell size/age and accounts for the background, SNR quantitative estimate is more appropriate for unbiased quantification of intracellular fluorescence over time. Ssb-Ypet, SsbF-mCh and mCh-ParB foci were detected using MicrobeJ Maxima detection function, and foci localisation and fluorescence intensity were extracted and plotted automatically. Plots presenting time lapse data were either aligned to the first frame where the transconjugant cell exhibits a conjugative Ssb-Ypet focus (ssDNA acquisition) or a mCh-ParB focus (ss-to-dsDNA conversion) as indicated in the corresponding figure legend.

### Statistical analysis

P-value significance was analysed running specific statistical tests on the GraphPad Prism software. Single-cell data from quantitative microscopy analysis were extracted from the MicrobeJ interface and transferred to GraphPad. P-value significance of single-cell quantitative data was performed using unpaired non-parametric Mann-Whitney statistical test, which allows to compare differences between independent data groups without normal distribution assumption. P-value significance for the frequency of transconjugants obtained by plating assays were evaluated using One-way analysis of variance (ANOVA) with Dunnetts multiple comparisons test, which allows to determine the statistical significance of differences observed between the means of three or more independent experimental groups against a control group mean (corresponding to the F*wt*). When required, P-value and significance are indicated on the figure panels and within the corresponding legend.

## Supporting information

MovieS1 ssDNA plasmid self-transfer

MovieS2 plasmid self-transfer triggers cell filamentation

MovieS3 bidirectionnal plasmid self-transfer

## Acknowledgments

We thank Pauline Rouzé and Pierre Bogaert for valuable help with sequencing of suppressor plasmids. We thank Sarah Bigot for valuable input during the project implementation

## Funding

This work was supported by funding from the Foundation for Medical Research, Grant number FRM-EQU202103012587 to C.L. and A.C.) and from the French National Research Agency (Grant numbers ANR-22-CE12-0032 and ANR-23-CE12-0037).

## Author contributions

C.L. provided funding. C.L. and A.C. conceptualized the study. A.C. and N.F. performed experiments and analyzed the data. C.L. and A.C. wrote the paper and prepared the figures, with input from N.F.

## Declaration of interests

The authors declare no competing interests.

## Notes

### Competing Interest Statement

The authors have declared no competing interest.

