## Supplemental File for "Exclusion Systems Preserve Host Cell Homeostasis and fitness, Ensuring Successful Dissemination of Conjugative Plasmids and Associated Resistance Genes"

This file includes:

Figures S1 to S5
Tables S1 to S3

Other Supplementary Information for this manuscript include the following:

Movies S1 to S3

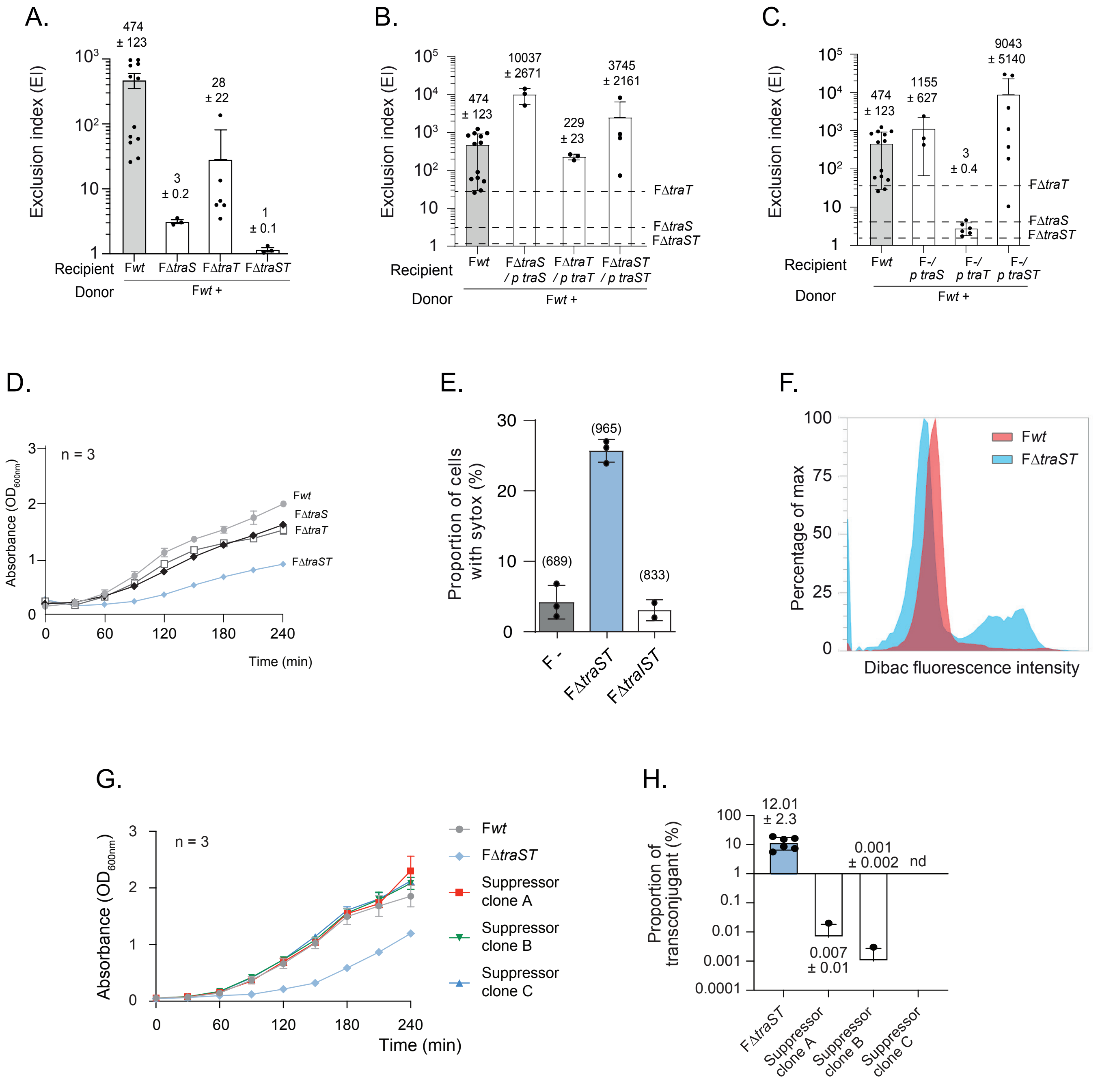
A.
B.
C.
D.
E.
F.
G.
H.
Exclusion index (EI)
Recipient
Donor
Fwt +
Absorbance (OD600nm)
Time (min)
Proportion of cells with sytox (%)
Percentage of max
Dibac fluorescence intensity
Proportion of transconjugant (%)
Suppressor clone A
Suppressor clone B
Suppressor clone C
n = 3
nd

Figure S1

Figure S1. Consequences of exclusion absence result in uncontrolled self-transfer

(A) Exclusion levels of recipient cells during F*wt* transfer from wild-type donors to recipient cells with compromised exclusion mechanisms due to *ΔtraS, ΔtraT* and *ΔtraST* deletions. The mean and SD are calculated from at least three biological replicates (black dots). (B) Histogram showing the exclusion levels of recipient cells during F*wt* transfer from wild-type donors to recipient cells carrying mutant plasmids, complemented with exclusion proteins. The exclusion levels of non-complemented mutants are indicated by a dashed line. The mean and SD are calculated from at least three biological replicates (black dots). (C) Histogram showing the exclusion levels of recipient cells during F*wt* transfer from wild-type donors to plasmid-free recipient cells complemented with exclusion proteins. The exclusion levels of non-complemented mutants are indicated by a dashed line. The mean and SD are calculated from at least three biological replicates (black dots). (D) Growth curves of F*wt,* FΔ*traS,* FΔ*traT* and FΔ*traST* mutant cells, measured by OD600 every 30 minutes over 240-minutes. Cells were grown in LB medium in triplicate, only the average curve is shown. (E) Proportion of cells with compromised plasma membranes, as determined by Sytox staining, in cells without a plasmid and in cells carrying FΔtraST or FΔtraIST plasmids. (F) Quantification of depolarized cells using DiBAC staining in cells carrying F*wt,* and F*ΔtraST.* (G) Growth curves of F*wt,* FΔ*traST* and suppressor clones, measured by OD600 every 30 minutes over 240-minutes. Cells were grown in LB medium in triplicate, only the average curve is shown. (H) Proportion of transconjugants obtained from suppressor plasmids (clone A, B and C) transferred into plasmid-free *wt* recipient cells. Bars represent the mean and SD from at least three independent biological replicates (black dots).

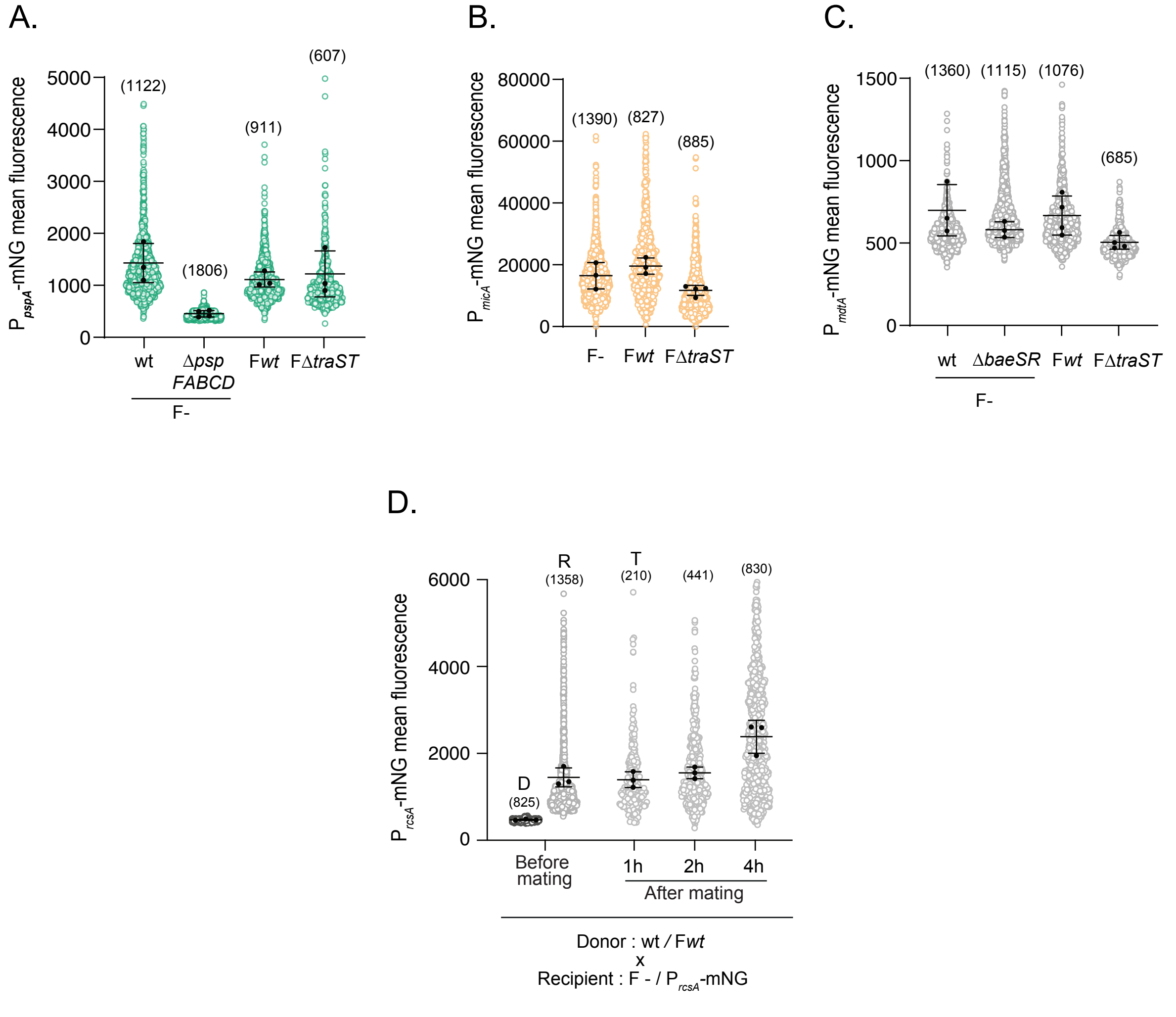
A.
$P_{pspA}$-mNG mean fluorescence
5000
4000
3000
2000
1000
0
(1122)
(1806)
(911)
(607)
wt
Δ*psp FABCD*
F*wt*
FΔ*traST*
F-
B.
$P_{micA}$-mNG mean fluorescence
80000
60000
40000
20000
0
(1390)
(827)
(885)
F-
F*wt*
FΔ*traST*
C.
$P_{mdtA}$-mNG mean fluorescence
1500
1000
500
0
(1360)
(1115)
(1076)
(685)
wt
Δ*baeSR*
F*wt*
FΔ*traST*
F-
D.
$P_{rcsA}$-mNG mean fluorescence
6000
4000
2000
0
D
(825)
R
(1358)
T
(210)
(441)
(830)
Before mating
1h
2h
4h
After mating
Donor : wt / F*wt*
x
Recipient : F - / $P_{rcsA}$-mNG

Figure S2

Figure S2. Membrane stress induced by deregulated transfer

**(A-C)** Quantification of envelope stress response (ESR) induction using different transcriptional reporters measured by fluorescence microscopy. **(A)** $P_{pspA}$-mNG **(B)** $P_{micA}$-mNG, and **(C)** $P_{mdtA}$-mNG were monitored in plasmid free strain (F-), F*wt*, F*ΔtraST* and ESR-deficient mutant strains (Δ*pspFABCD* and Δ*baeSR).* Each dot represents an individual cell; black dots indicate the mean fluorescence value for each independent biological replicate. **(D)** Quantification of $P_{rcsA}$-mNG reporter induction at the single-cell level before mating and at 1, 2 and 4 hours of mating, in cells that have acquired F*wt* plasmid (transconjugant).

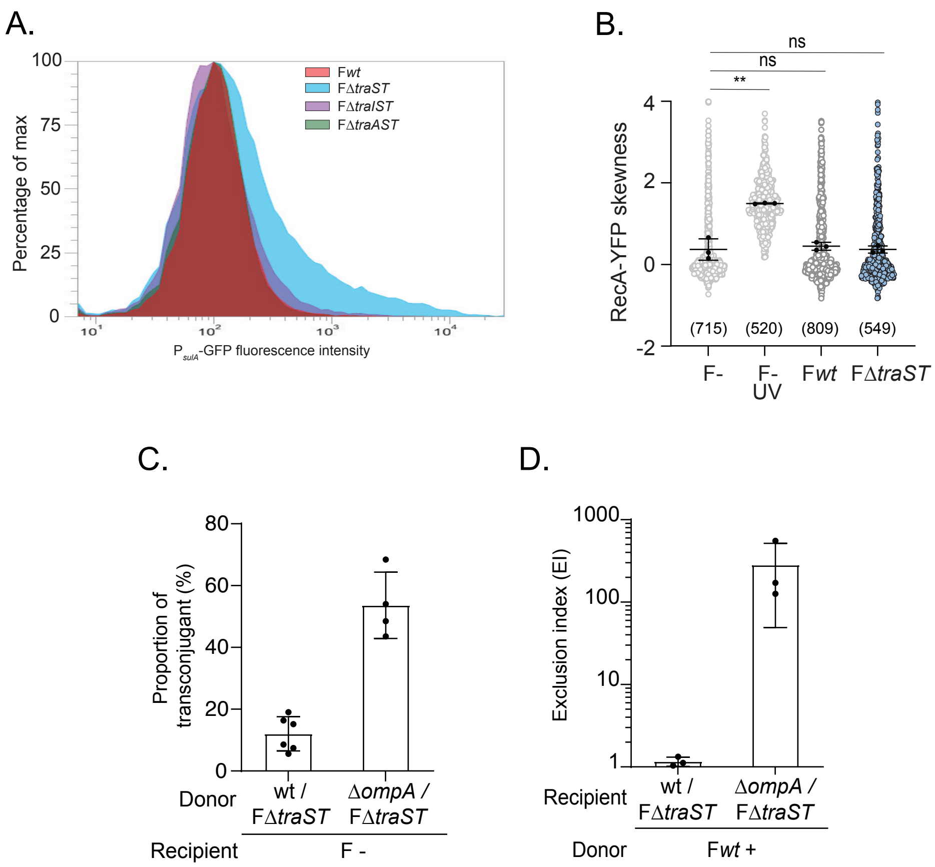
A.
B.
C.
D.
Percentage of max
Fwt
FΔtraST
FΔtraIST
FΔtraAST
P sulA-GFP fluorescence intensity
RecA-YFP skewness
ns
ns
**
(715) (520) (809) (549)
F-
F- UV
Fwt
FΔtraST
Proportion of transconjugant (%)
Donor
wt / FΔtraST
ΔompA / FΔtraST
Recipient
F -
Exclusion index (EI)
Recipient
wt / FΔtraST
ΔompA / FΔtraST
Donor
Fwt +

Figure S3

Figure S3. Analysis of the SOS response induced by deregulated transfer

**(A)** Quantification of SOS response induction using $P_{sulA}$-GFP reporter measured by flow-cytometry in F*wt,* F*ΔtraST,* F*ΔtraIST* and F*ΔtraAST* strains. **(B)** Jitter plot showing the skewness of RecA-YFP fluorescence in plasmid-free cells (F-), with and without UV treatment, as well as in F*wt* and F*ΔtraST* cells. Each dot represents an individual cell; black dots indicate the mean fluorescence value for each independent biological replicate. Statistical significance was assessed using an unpaired t-test on replicate means (***P* < 0.005). **(C)** Proportion of transconjugants obtained from F*ΔtraST* transfer from *wt* and *ΔompA* donors to plasmid-free *wt* recipient cells. The mean and SD are calculated from at least three biological replicates (black dots). **(D)** Exclusion level of recipient cells during F*wt* transfer from wild-type donors to *wt* and *ΔompA* recipient cells, both carrying F*ΔtraST* plasmid. The mean and SD are calculated from at least three biological replicates (black dots).

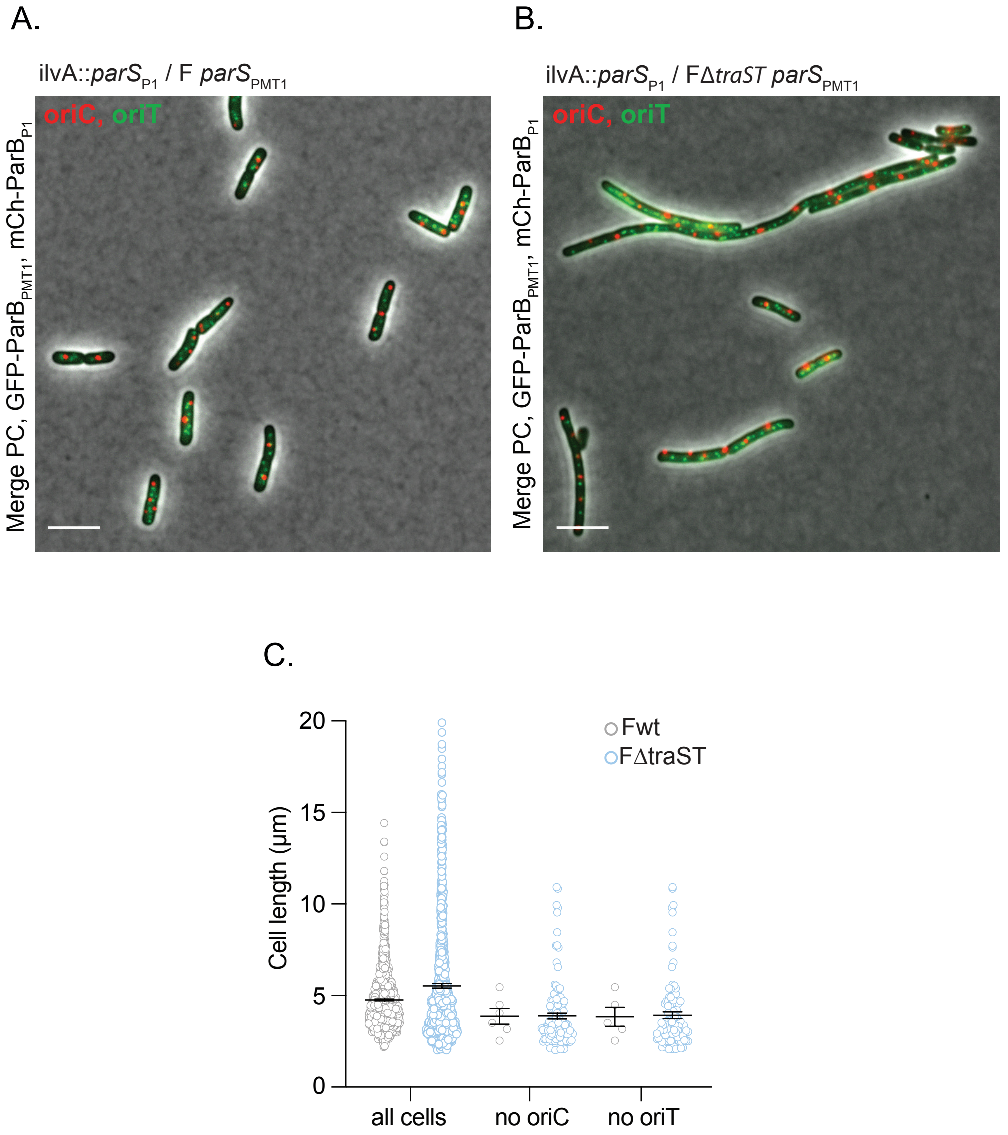
A.
ilvA::parSP1 / F parSPMT1
oriC, oriT
Merge PC, GFP-ParBPMT1, mCh-ParBP1
B.
ilvA::parSP1 / FΔtraST parSPMT1
oriC, oriT
Merge PC, GFP-ParBPMT1, mCh-ParBP1
C.
Cell length (µm)
0
5
10
15
20
Fwt
FΔtraST
all cells
no oriC
no oriT

Figure S4

Figure S4 : Effect of exclusion loss on plasmid stability

A. Representative fluorescence microscopy images of clonal population cells carrying a chromosomal insertion of *parS*$_{P1}$ at the *ilvA* locus, near the origin of replication (*oriC*), and harboring the F*wt* plasmid with a *parS*$_{PMT1}$ insertion adjacent to the origin of transfer (*oriT*). B. Representative fluorescence microscopy images of clonal population cells carrying a chromosomal insertion of *parS*$_{P1}$ at the *ilvA* locus, near the origin of replication (*oriC*), and harboring the FΔ*traST* plasmid with a *parS*$_{PMT1}$ insertion adjacent to the origin of transfer (*oriT*). C. Histogram showing the distribution of cell lengths for F*wt* and FΔ*traST* cells, compiled from at least three independent experiments. D. Scatter plot of cell lengths for F*wt* and FΔ*traST* cells, shown for the entire population, cells lacking *oriC*, and cells lacking *oriT*.

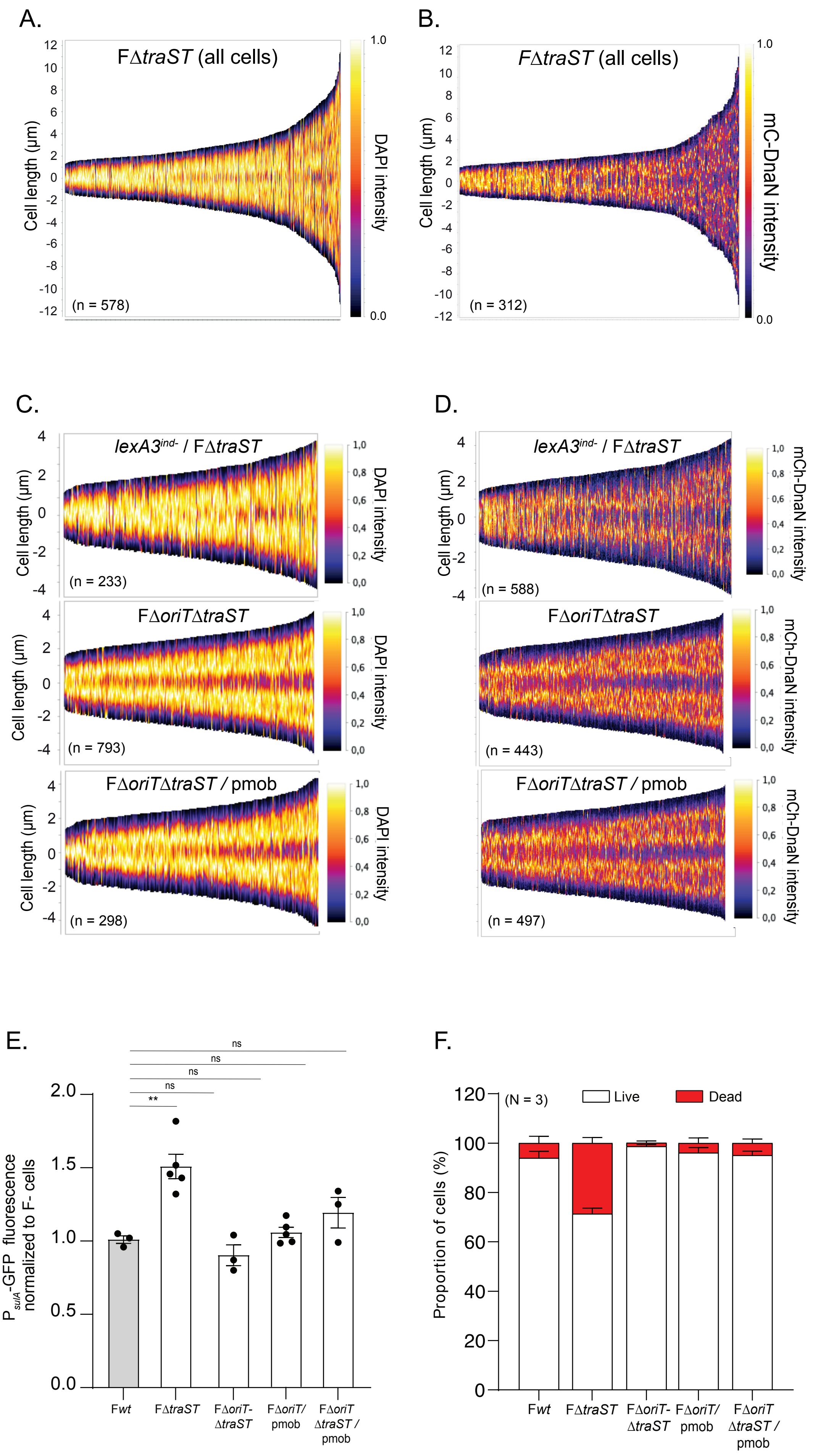
A.
FΔtraST (all cells)
Cell length (µm)
DAPI intensity
(n = 578)
B.
FΔtraST (all cells)
mC-DnaN intensity
(n = 312)
C.
lexA3ind- / FΔtraST
(n = 233)
FΔoriTΔtraST
(n = 793)
FΔoriTΔtraST / pmob
(n = 298)
D.
lexA3ind- / FΔtraST
mCh-DnaN intensity
(n = 588)
FΔoriTΔtraST
(n = 443)
FΔoriTΔtraST / pmob
(n = 497)
E.
PsulA-GFP fluorescence normalized to F- cells
ns
**
Fwt
FΔtraST
FΔoriT-ΔtraST
FΔoriT/ pmob
FΔoriT ΔtraST / pmob
F.
(N = 3)
Live
Dead
Proportion of cells (%)
Fwt
FΔtraST
FΔoriT-ΔtraST
FΔoriT/ pmob
FΔoriT ΔtraST / pmob

Figure S5

Figure S5 : Unregulated self-transfer induces cell cycle-disruption

**(A)** Demographs showing the distribution of DAPI staining across the full population of FΔtraST-carrying cells, with cell lengths ranging from 4 to 24 µm. The number of cells analyzed (n) is indicated, based on data from three independent experiments. **(B)** Localization of the mCh-DnaN fusion throughout the cell cycle, across the entire population, with cell lengths ranging from 4 to 24 µm. The number of cells analyzed (n) is indicated, based on data from three independent experiments. **(C)** Nucleoid positioning visualized by DAPI staining and **(D)** replisome localization using a mCh-DnaN fusion in *lexA3*$^{ind-}$ strains carrying F*ΔtraST* and in wt strains carrying F*ΔoriTΔtraST* with or without pmob plasmid. For both panels, demographs show signal distribution in cells sorted by length (4–8 µm). **(F)** Quantification of SOS response induction using $P_{sulA}$-GFP reporter in strains carrying F*wt,* F*ΔtraST,* and F*ΔoriTΔtraST* with or without pmob plasmid. Fluorescence intensity was measured by fluorescence microscopy and normalized to plasmid-free (F-) cells. Bars represent the mean ± standard deviation (SD) from at least three independent experiments. **(G)** Histograms showing the proportion of live and dead in F*wt,* F*ΔtraST,* and F*ΔoriTΔtraST* with or without pmob plasmid as determined by the Live/Dead staining assay. Bars represent mean and standard deviation (SD) from at least three independent experiments.

| Strain | Relevant genotype[a, b, c] | Source or reference[a] |
|---|---|---|
| **MG1655 and derivatives** | | |
| LY5 | MS388 / F-Tn*10*, $parS_{P1}$-*FRT* | Nolivos *et al.,* 2019 |
| LY117 | MS388 *ssb-ypet-FRT-kan-FRT* | Nolivos *et al.,* 2019 |
| LY128 | MS388 *ssb-ypet-FRT* | Nolivos *et al.,* 2019 |
| LY318 | MS388 / pSN70 | Nolivos *et al.,* 2019 |
| LY583 | MS388 Δ*ompA :: FRT-kan-FRT* | MS388 x P1.LY575 to $Kan^r$ |
| LY597 | MS388 Δ*ompA :: FRT* | Derivative of LY583, *kan* removed via pCP20 |
| LY701 | MS388 *dnaN-mCherry-FRT-kan-FRT* | MS388 x P1. *FRT-kan-FRT-mCh-dnaN* to $Kan^r$ |
| LY832 | MS388 *ilva :: Ery* | MS388 x P1.Erythromycin to $Ery^r$ |
| LY833 | MS388 / F-Tn*10*, $parS_{P1}$-*FRT,* Δ*oriT::FRT-kan-FRT* | LY5 x P1.LY820 to $Kan^r$ |
| LY873 | MS388 / F-Tn*10*, $parS_{P1}$-*FRT,* Δ*oriT::FRT* | Derivative of LY833, *kan* removed via pCP20 |
| LY889 | MS388 / F-Tn*10*, $parS_{P1}$-*FRT,* Δ*oriT::FRT-kan-FRT /* pSEVA-*oriT* | pSEVA-*oriT* x LY873 to $Ap^r$ |
| LY1107 | MS388 $lexA3^{ind-}$ | MS388 x P1 $lexA3^{ind-}$to $Tc^r$ |
| LY1553 | MS388 / F-Tn*10*, *repE :: pBiofab-sfgfp-FRT-kan-FRT* | Conjugation LY1552 x MS388 $St^r$ $Kan^r$ |
| LY1650 | MS428 / F-Tn*10*, *repE :: pTac-mcherry-FRT-kan-FRT* | Conjugation LY1642 x MS428 to $LAC^-$ $St^r$ $Tc^r$ |
| LY1651 | MS388 Δ*lacZ* | MS388 x *lacZ* deletion |
| LY1773 | MS428 / F-Tn*10*, *repE :: pBiofab-sfgfp-FRT-kan-FRT* | Conjugation LY1553 x MS428 to $LAC^-$ $St^r$ $Tc^r$ |
| LY1784 | MS388 *ilva :: pTac-mCherry* / F-Tn*10*, *repE :: pBiofab-sfgfp-FRT-kan-FRT* | Conjugation LY1773 x LY1593 to $St^r$ $Tc^r$ |
| LY1787 | MS388 *ilva :: Ery* / F-Tn*10*, $parS_{PMT1}$-*FRT,* Δ*traS::FRT-cat-FRT* | Conjugation LY1774 x LY832 to $Ery^r$ $Tc^r$ |
| LY1794 | MS388 *ilva :: Ery* / F-Tn*10*, $parS_{PMT1}$-*FRT,* Δ*traST::FRT-cat-FRT* | Conjugation LY1775 x LY832 to $Ery^r$ $Tc^r$ |
| LY1808 | MS388 *ilva :: Ery* / F-Tn*10*, $parS_{PMT1}$-*FRT-cat-FRT* | Conjugation LY161 x LY832 to $Ery^r$ $Tc^r$ |

| | | |
|---|---|---|
| LY1809 | MS388 *ilva :: Ery* / F-Tn*10*, *parS*$_{PMT1}$*-FRT, ΔtraT::FRT-cat-FRT* | Conjugation LY1801 x LY832 to Ery$^r$ Tc$^r$ |
| LY1818 | MS388 *ilva :: Ery* / F-Tn*10*, *parS*$_{PMT1}$*-FRT, ΔtraS::FRT-cat-FRT* / pAC2 | pAC2 x LY1787 to Ap$^r$ |
| LY1821 | MS388 *ilva :: Ery* / F-Tn*10*, *parS*$_{PMT1}$*-FRT, ΔtraST::FRT-cat-FRT* / pAC4 | pAC4 x LY1794 to Ap$^r$ |
| LY1828 | MS388 *ilva :: Ery* / pAC2 | pAC2 x LY832 to Ap$^r$ |
| LY1829 | MS388 *ilva :: Ery* / pAC3 | pAC3 x LY832 to Ap$^r$ |
| LY1830 | MS388 *ilva :: Ery* / pAC4 | pAC4 x LY832 to Ap$^r$ |
| LY1836 | MS388 *ilva :: Ery* / F-Tn*10*, *parS*$_{PMT1}$*-FRT, ΔtraT::FRT-cat-FRT* / pAC3 | pAC3 x LY1809 to Ap$^r$ |
| LY1880 | MS388 *ssb-ypet-FRT* / F-Tn*10*, *parS*$_{PMT1}$*-FRT-cat-FRT* | Conjugation LY161 x LY318 to St$^r$ Tc$^r$ |
| LY1883 | MS388 *ssb-ypet-FRT* / F-Tn*10*, *parS*$_{PMT1}$*-FRT, ΔtraST::FRT-cat-FRT* | Conjugation LY1775 x LY128 to St$^r$ Tc$^r$ |
| LY2159 | MS388 *ilvA :: parS*$_{P1}$*-FRT* / F-Tn*10*, *parS*$_{PMT1}$*-FRT-cat-FRT* | Conjugation LY161 x LY196 to St$^r$ Tc$^r$ |
| LY2160 | MS388 *ilvA :: parS*$_{P1}$*-FRT* / F-Tn*10*, *parS*$_{PMT1}$*-FRT, ΔtraS::FRT-cat-FRT* | Conjugation LY1774 x LY196 to St$^r$ Tc$^r$ |
| LY2161 | MS388 *ilvA :: parS*$_{P1}$*-FRT* / F-Tn*10*, *parS*$_{PMT1}$*-FRT, ΔtraT::FRT-cat-FRT* | Conjugation LY1801 x LY196 to St$^r$ Tc$^r$ |
| LY2162 | MS388 *ilvA :: parS*$_{P1}$*-FRT* / F-Tn*10*, *parS*$_{PMT1}$*-FRT, ΔtraST::FRT-cat-FRT* | Conjugation LY161 x LY1775 to St$^r$ Tc$^r$ |
| LY2168 | MS388 *ilvA :: parS*$_{P1}$*-FRT* / F-Tn*10*, *parS*$_{PMT1}$*-FRT-cat-FRT* / p2973 | p2973 x LY2159 to Ap$^r$ |
| LY2169 | MS388 *ilvA :: parS*$_{P1}$*-FRT* / F-Tn*10*, *parS*$_{PMT1}$*-FRT, ΔtraS::FRT-cat-FRT* / p2973 | p2973 x LY2160 to Ap$^r$ |
| LY2170 | MS388 *ilvA :: parS*$_{P1}$*-FRT* / F-Tn*10*, *parS*$_{PMT1}$*-FRT, ΔtraT::FRT-cat-FRT* / p2973 | p2973 x LY2161 to Ap$^r$ |
| LY2171 | MS388 *ilvA :: parS*$_{P1}$*-FRT* / F-Tn*10*, *parS*$_{PMT1}$*-FRT, ΔtraST::FRT-cat-FRT* / p2973 | p2973 x LY2162 to Ap$^r$ |
| LY2277 | MS388 Δ*lacZ* / F-Tn*10*, *parS*$_{PMT1}$*-FRT-cat-FRT* | Conjugation LY1808 x LY1651 LAC$^-$ St$^r$ Tc$^r$ |

| | | |
|---|---|---|
| LY2280 | MS388 Δ*lacZ* / F-Tn*10*, *parS*$_{PMT1}$*-FRT, ΔtraST::FRT-cat-FRT* | Conjugation LY1794 x LY1651 LAC$^{-}$ St$^{r}$ Tc$^{r}$ |
| LY2330 | MS388 *ilva :: Ery* / F-Tn*10*, *parS*$_{PMT1}$*-FRT, ΔtraI::FRT-kan-FRT, ΔtraST::FRT-cat-FRT* | Conjugation LY2342 x LY832 to Ery$^{r}$ Tc$^{r}$ |
| LY2494 | MS388 *lexA3*$^{ind-}$ / F-Tn*10*, *parS*$_{PMT1}$*-FRT-cat-FRT* | Conjugation LY161 x LY1107 to St$^{r}$ Tc$^{r}$ |
| LY2496 | MS388 *lexA3*$^{ind-}$ / F-Tn*10*, *parS*$_{PMT1}$*-FRT, ΔtraST::FRT-cat-FRT* | Conjugation LY2280 x LY1107 to Tc$^{r}$ Cm$^{r}$ |
| LY2584 | MS388 *ilvA :: Ery* / F-Tn*10*, *parS*$_{PMT1}$*-FRT-repE ::pBiofab-sfgfp-kan-FRT, ΔtraST::FRT-cat-FRT* | LY1794 x P1.LY1553 to St$^{r}$ Kan$^{r}$ |
| LY2585 | MS388 *ilvA :: Ery* / F-Tn*10*, *parS*$_{PMT1}$*-FRT-repE ::ptac-mcherry-kan-FRT, ΔtraST::FRT-cat-FRT* | LY1794 x P1.LY1650 to St$^{r}$ Kan$^{r}$ |
| LY2620 | MS388 Δ*ompA* / F-Tn*10*, *parS*$_{PMT1}$*-FRT, ΔtraST::FRT-cat-FRT* | Conjugation LY1775 x LY597 to St$^{r}$ Tc$^{r}$ |
| LY2652 | MS388 / F-Tn*10*, *parS*$_{P1}$*-FRT, ΔoriT::FRT-kan-FRT, ΔtraST::FRT-cat-FRT* | LY1794 x P1.LY833 to Kan$^{r}$ Cm$^{r}$ |
| LY2656 | MS388 / F-Tn*10*, *parS*$_{P1}$*-FRT, ΔtraST::FRT-cat-FRT* | Conjugation LY2697 x MS388 to St$^{r}$ Tc$^{r}$ |
| LY2660 | MS388 *ilva :: Ery* / F-Tn*10*, *parS*$_{PMT1}$*-FRT, ΔtraST::FRT-cat-FRT* / p2973 | p2973 x LY1787 to Ap$^{r}$ |
| LY2661 | MS388 / F-Tn*10*, *parS*$_{P1}$*-FRT, ΔtraST::FRT-cat-FRT* / p2973 | p2973 x LY2656 to Ap$^{r}$ |
| LY2664 | MS388 / F-Tn*10*, *parS*$_{PMT1}$*-FRT, ΔoriT::FRT, ΔtraST::FRT* | Derivative of LY2652, *kan* and *cat* removed via pCP20 |
| LY2674 | MS388 / F-Tn*10*, *parS*$_{P1}$*-FRT* | Derivative of LY2663 *cat* removed via pCP20 |
| LY2678 | MS388 / F-Tn*10*, *parS*$_{P1}$*-FRT, ΔoriT::FRT, ΔtraST::FRT* / pSEVA-*oriT* | pSEVA-*oriT* x LY2664 to Ap$^{r}$ |
| LY2872 | MS388, *recA-recA-yfp-FRT-kan-FRT* | MS388 x P1.LY2853 to St$^{r}$ Kan$^{r}$ |
| LY2873 | MS388, *recA-recA-yfp-FRT-kan-FRT / -*Tn*10*, *parS*$_{PMT1}$*-FRT-cat-FRT* | Conjugation LY2277 x LY2872 to St$^{r}$ Kan$^{r}$ Tc$^{r}$ Cm$^{r}$ |
| LY2877 | MS388, *recA-recA-yfp-FRT-kan-FRT* / F-Tn*10*, *parS*$_{PMT1}$*-FRT, ΔtraS::FRT-cat-FRT* | Conjugation LY2280 x LY2872 to St$^{r}$ Kan$^{r}$ Tc$^{r}$ Cm$^{r}$ |

| | | |
|---|---|---|
| LY2975 | MG1655 CF1648 Δ*baeSR* :: *FRT- kan-FRT* | Rousseau *et al.,* 2023 |
| LY2976 | MG1655 CF1648 ΔcpxQPRA :: *FRT-kan-FRT* | Rousseau *et al.,* 2023 |
| LY2977 | MG1655 CF1648 Δ*pspFABCD* :: *FRT-kan-FRT* | Rousseau *et al.,* 2023 |
| LY2978 | MG1655 CF1648 ΔrcsDB :: *FRT- kan-FRT* | Rousseau *et al.,* 2023 |
| LY2989 | MS388 Δ*baeSR* :: *FRT- kan-FRT* | MS388 x P1.LY2975 to $St^r$ $Kan^r$ |
| LY2991 | MS388 Δ*pspFABCD* :: *FRT- kan-FRT* | MS388 x P1.LY2977 to $St^r$ $Kan^r$ |
| LY3071 | MS388 *ilvA :: Ery* / F-Tn*10*, *parS*$_{PMT1}$-*FRT-cat-FRT* / pAC13 | pAC13 x LY1808 to $Gm^r$ |
| LY3073 | MS388 / F-Tn*10*, *parS*$_{PMT1}$*-FRT,* Δ*traST::FRT-cat-FRT* / pAC13 | pAC13 x LY1794 to $Gm^r$ |
| LY3075 | MS388 *mcherry-dnaN* / F-Tn*10*, *parS*$_{PMT1}$*-FRT-cat-FRT* | Conjugation LY1808 x LY701 to $St^r$ $Kan^r$ $Tc^r$ $Cm^r$ |
| LY3075 | MS388 *mcherry-dnaN* / F-Tn*10*, *parS*$_{PMT1}$*-FRT,* Δ*traST::FRT-cat-FRT* | Conjugation LY1794 x LY701 to $St^r$ $Kan^r$ $Tc^r$ $Cm^r$ |
| LY3078 | MS388 *ilvA :: Ery* / F-Tn*10*, *parS*$_{PMT1}$-*FRT-cat-FRT* / pAC14 | pAC14 x LY1808 to $Gm^r$ |
| LY3079 | MS388 *ilvA :: Ery* / F-Tn*10*, *parS*$_{PMT1}$-*FRT-cat-FRT* / pAC16 | pAC16 x LY1808 to $Gm^r$ |
| LY3080 | MS388 / F-Tn*10*, *parS*$_{PMT1}$*-FRT,* Δ*traST::FRT-cat-FRT* / pAC14 | pAC14 x LY1794 to $Gm^r$ |
| LY3081 | MS388 / F-Tn*10*, *parS*$_{PMT1}$*-FRT,* Δ*traST::FRT-cat-FRT* / pAC16 | pAC16 x LY1794 to $Gm^r$ |
| LY3086 | MS388 / pAC13 | pAC13 x MS388 to $Gm^r$ |
| LY3087 | MS388 / pAC14 | pAC14 x MS388 to $Gm^r$ |
| LY3089 | MS388 / pAC16 | pAC16 x MS388 to $Gm^r$ |
| LY3091 | MS388 Δ*baeSR* :: *FRT- kan-FRT* / pAC16 | pAC16 x LY2989 to $Gm^r$ |
| LY3093 | MS388 Δ*pspFABCD* :: *FRT- kan-FRT* / pAC13 | pAC13 x LY2989 to $Gm^r$ |
| LY3387 | MGT Δ*rcsDB :: FRT- kan-FRT* | MGT x P1.LY2978 to $Kan^r$ |

| | | |
|---|---|---|
| LY3388 | MGT Δ*cpxQPRA :: FRT- kan-FRT* | MGT x P1.LY2976 to Kan$^r$ |
| LY3399 | MGT / F-Tn*10*, *parS*$_{PMT1}$*-FRT* | Conjugation LY162 x MGT to St$^r$ Tc$^r$ |
| LY3400 | MGT / F-Tn*10*, *parS*$_{PMT1}$*-FRT-cat-FRT* | Conjugation LY161 x MGT to St$^r$ Tc$^r$ Cm$^r$ |
| LY3401 | MGT / F-Tn*10*, *parS*$_{PMT1}$*-FRT,* Δ*traS::FRT-cat-FRT* | Conjugation LY1774 x MGT to St$^r$ Tc$^r$ Cm$^r$ |
| LY3402 | MGT / F-Tn*10*, *parS*$_{PMT1}$*-FRT,* Δ*traT::FRT-cat-FRT* | Conjugation LY1801 x MGT to St$^r$ Tc$^r$ Cm$^r$ |
| LY3403 | MGT / F-Tn*10*, *parS*$_{PMT1}$*-FRT,* Δ*traST::FRT-cat-FRT* | Conjugation LY1775 x MGT to St$^r$ Tc$^r$ Cm$^r$ |
| LY3404 | MGT / F-Tn*10*, *parS*$_{PMT1}$*-FRT,* Δ*traI::FRT-kan-FRT,* Δ*traST::FRT-cat-FRT* | Conjugation LY2342 x MGT to St$^r$ Tc$^r$ Cm$^r$ Kan$^r$ |
| LY3424 | MGT Δ*rcsDB :: FRT- kan-FRT /* F-Tn*10*, *parS*$_{PMT1}$*-FRT,* Δ*traST::FRT-cat-FRT* | Conjugation LY3403 x LY3387 to Tc$^r$ Cm$^r$ |
| LY3425 | MGT Δ*cpsQPRA :: FRT- kan-FRT /* F-Tn*10*, *parS*$_{PMT1}$*-FRT,* Δ*traST::FRT-cat-FRT* | Conjugation LY3403 x LY3388 to Tc$^r$ Cm$^r$ |
| LY3434 | MGT / F-Tn*10*, *parS*$_{PMT1}$*-FRT,* Δ*traA::FRT-kan-FRT,* Δ*traST::FRT-cat-FRT* | LY3399 x P1.LY2299 to Cm$^r$ Kan$^r$ |
| LY3670 | MGT / F-Tn*10*, *parS*$_{PMT1}$*-FRT-cat-FRT* / pUA66-P$_{sulA}$-GFP | pUA66-P$_{sulA}$-GFP x LY3400 to Gm$^r$ |
| LY3673 | MGT / F-Tn*10*, *parS*$_{PMT1}$*-FRT,* Δ*traST::FRT-cat-FRT* / pUA66-P$_{sulA}$-GFP | pUA66-P$_{sulA}$-GFP x LY3403 to Gm$^r$ |
| LY3679 | MGT / F-Tn*10*, *parS*$_{PMT1}$*-FRT,* Δ*traA::FRT-kan-FRT,* Δ*traST::FRT-cat-FRT* / pUA66-P$_{sulA}$-GFP | pUA66-P$_{sulA}$-GFP x LY3434 to Gm$^r$ |
| LY3686 | MGT / pUA66-P$_{sulA}$-GFP | pUA66-P$_{sulA}$-GFP x MGT to Gm$^r$ |
| LY3687 | MGT / F-Tn*10*, *parS*$_{PMT1}$*-FRT,* Δ*traI::FRT-kan-FRT,* Δ*traST::FRT-cat-FRT* / pUA66-P$_{sulA}$-GFP | pUA66-P$_{sulA}$-GFP x LY3404 to Gm$^r$ |
| LY3712 | MGT / pAC17 | pAC17 x LY3372 to Gm$^r$ |
| LY3713 | MGT / F-Tn*10*, *parS*$_{PMT1}$*-FRT-cat-FRT* / pAC15 | pAC15 x LY3400 to Gm$^r$ |
| LY3714 | MGT / F-Tn*10*, *parS*$_{PMT1}$*-FRT-cat-FRT* / pAC17 | pAC17 x LY3400 to Gm$^r$ |

| | | |
|---|---|---|
| LY3715 | MGT / F-Tn*10*, *parS*$_{PMT1}$*-FRT, ΔtraST::FRT-cat-FRT* / pAC15 | pAC15 x LY3403 to $Gm^r$ |
| LY3716 | MGT / F-Tn*10*, *parS*$_{PMT1}$*-FRT, ΔtraST::FRT-cat-FRT* / pAC17 | pAC17 x LY3403 to $Gm^r$ |
| LY3717 | MGT / F-Tn*10*, *parS*$_{PMT1}$*-FRT, ΔtraI::FRT-kan-FRT, ΔtraST::FRT-cat-FRT* / pAC15 | pAC15 x LY3404 to $Gm^r$ |
| LY3718 | MGT / F-Tn*10*, *parS*$_{PMT1}$*-FRT, ΔtraI::FRT-kan-FRT, ΔtraST::FRT-cat-FRT* / pAC17 | pAC17 x LY3404 to $Gm^r$ |
| LY3719 | MGT / F-Tn*10*, *parS*$_{PMT1}$*-FRT, ΔtraA::FRT-kan-FRT, ΔtraST::FRT-cat-FRT* / pAC15 | pAC15 x LY3434 to $Gm^r$ |
| LY3720 | MGT / F-Tn*10*, *parS*$_{PMT1}$*-FRT, ΔtraA::FRT-kan-FRT, ΔtraST::FRT-cat-FRT* / pAC17 | pAC17 x LY3434 to $Gm^r$ |
| LY3784 | MGT Δ*cpxQPRA :: FRT* | Derivative of LY3388, *kan* removed via pCP20 |
| LY3792 | MS388 *dnaN-mCherry* / F-Tn*10*, *parS*$_{PMT1}$*-FRT, ΔtraI::FRT-kan-FRT, ΔtraST::FRT-cat-FRT* | Conjugation LY2342 x LY701 to $Tc^r$ $Cm^r$ |
| LY3793 | MS388, *lexA3*$^{ind-}$ *dnaN-mCherry* / F-Tn*10*, *parS*$_{PMT1}$*-FRT, ΔtraST::FRT-cat-FRT* | LY2496 x P1.LY701 to $Cm^r$, $Kan^r$ |
| LY3811 | MGT Δ*cpxQPRA :: FRT* Δ*rcsDB ::FRT-kan-FRT* | LY3784 x P1.LY3387 x to $Kan^r$ |
| LY3812 | MGT Δ*cpxQPRA :: FRT* Δ*rcsDB ::FRT-kan-FRT* | Conjugation LY3403 x LY3811 to $Tc^r$, $Cm^r$ |
| LY3832 | MGT / F-Tn*10*, *parS*$_{PMT1}$*-FRT, ΔtraS::FRT-cat-FRT* / pAC15 | pAC15 x LY3401 to $Gm^r$ |
| LY3833 | MGT / F-Tn*10*, *parS*$_{PMT1}$*-FRT, ΔtraS::FRT-cat-FRT* / pAC17 | pAC17 x LY3401 to $Gm^r$ |
| LY3835 | MGT / F-Tn*10*, *parS*$_{PMT1}$*-FRT, ΔtraT::FRT-cat-FRT* / pAC15 | pAC15 x LY3402 to $Gm^r$ |
| LY3836 | MGT / F-Tn*10*, *parS*$_{PMT1}$*-FRT, ΔtraT::FRT-cat-FRT* / pAC17 | pAC17 x LY3402 to $Gm^r$ |
| LY4134 | MS388 *FRT-kan-FRT-mcherry-dnaN* / F-Tn*10*, *parS*$_{PMT1}$*-FRT, ΔoriT-FRT, ΔtraST::FRT-cat-FRT* | LY2664 x P1.LY701 to $Cm^r$, $Kan^r$ |

| | | |
|---|---|---|
| LY4142 | MS388 *FRT-kan-FRT-mcherry-dnaN* / F-Tn*10*, *parS*$_{PMT1}$*-FRT, ΔoriT-FRT, ΔtraST::FRT-cat-FRT /* pSEVA-*oriT* | pSEVA-*oriT* x LY4134 to Ap$^r$ |
| MG1655 | λ$^-$, *rph-1* | Coli Genetic Stock Center (CGSC) #6300 |
| MS388 | MG1655 *rpsL* (St$^R$) | Gift from F. Cornet |
| Other genetic backgrounds | | |
| K603 | F$^+$ [F1-10(Tn*10*)], *thr-1, araC14, leuB6*(Am)*, lacY1, glnX44*(AS)*, galK2*(Oc)*, galT22, λ*$^-$*, ΔtrpE63, xylA5, mtl-1, thiE1* | Coli Genetic Stock Center CGSC #6451<br>Strain carrying the F-Tn*10* natural isolate with Tn*10* transposon in the intergenic region *ybdB-ybfA*. GenBank accession number MK492260) |
| DY330 | W3110 *ΔlacU169, gal490, λcI857, Δ(cro-bioA)* | Yu *et al.,* 2000 |
| LY161 | DY330 / F-Tn*10*, *parS*$_{PMT1}$*-FRT-cat-FRT* | Nolivos *et al.,* 2019 |
| LY162 | DY330 / F-Tn*10-parS*$_{PMT1}$*-FRT* | Nolivos *et al.,* 2019 |
| LY575 | BW25113 Δ*OmpA772 :: FRT-kan-FRT* | JW0940 KEIO collection |
| LY1552 | DY330 / F-Tn*10*, *repE :: pBiofab-sfgfp-FRT-kan-FRT* | λred *pBiofab-sfgfp-FRT-kan-FRT* construct at the F plasmid intergenic *repE locus* (OL688/OL117) |
| LY1577 | DY330 *ilvA :: ptac-mcherry-FRT-kan-FRT* | Λred *ptac-mCherry-FRT-kan-FRT* at the endogenous *ilvA locus* (OL743/OL383) |
| LY1642 | DY330 / F-Tn*10*, *repE :: pTac-mcherry-FRT-kan-FRT* | λred *pTac-mcherry-FRT-kan-FRT* construct at the F plasmid intergenic *repE locus* (OL817/OL117) |
| LY1774 | DY330 / F-Tn*10*, *parS*$_{PMT1}$*-FRT*, Δ*traS::FRT-cat-FRT* | λred Δ*traS :: cat* construct at the endogenous F plasmid *locus* (OL737/OL738) |
| LY1775 | DY330 / F-Tn*10*, *parS*$_{PMT1}$*-FRT*, Δ*traST::FRT-cat-FRT* | λred Δ*traST :: cat* construct at the endogenous F plasmid *locus* (OL737/OL734) |
| LY1801 | DY330 / F-Tn*10*, *parS*$_{PMT1}$*-FRT*, Δ*traT::FRT-cat-FRT* | λred Δ*traT :: cat* construct at the endogenous F plasmid *locus* (OL733/OL734) |
| LY2293 | DY330 / F-Tn*10*, *parS*$_{PMT1}$*-FRT*, Δ*traA::FRT-kan-FRT* | λred Δ*traA :: kan* construct at the endogenous F plasmid *locus* (OL768/OL767) |
| LY2295 | DY330 / F-Tn*10*, *parS*$_{PMT1}$*-FRT*, Δ*traI::FRT-kan-FRT* | λred Δ*traI :: kan* construct at the endogenous F plasmid *locus* (OL944/OL945) |

| | | |
|---|---|---|
| LY2299 | DY330 / F-Tn*10*, *parS$_{PMT1}$-FRT*, Δ*traA::FRT-kan-FRT,* Δ*traST::FRT-cat-FRT* | λred Δ*traST :: cat* construct at the endogenous F plasmid *locus* (OL737/OL734) on LY2293 |
| LY2337 | DY330 / F-Tn*10*, *parS$_{PMT1}$-FRT*, Δ*traI::FRT-kan-FRT,* Δ*traST::FRT-cat-FRT* | λred Δ*traST :: cat* construct at the endogenous F plasmid *locus* of LY2295 (OL737/OL734) |
| LY2342 | DY330 / F-Tn*10*, *parS$_{PMT1}$-FRT*, Δ*traI::FRT-kan-FRT,* Δ*traST::FRT-cat-FRT* / pTrc-traI | pTrc-traI x LY2337 to Ap$^r$ |
| LY2681 | DY330 / F-Tn*10-parS$_{P1}$-FRT* | Conjugation LY2674 x DY330 to LAC$^-$, Tc$^r$ |
| LY2697 | DY330 / F-Tn*10*, *parS$_{P1}$-FRT*, Δ*traST::FRT-cat-FRT* | λred Δ*traST :: cat* construct at the endogenous F plasmid *locus* (OL737/OL734) on LY2681 |

[a]The abbreviations *bla*, *kan Gm and cat* refer to insertions genes conferring resistance to ampicillin (Ap$^r$) kanamycin (Kn$^r$), gentamycine (Gm$^r$) and chloramphenicol (Cm$^r$); *rpsl* refers to spontaneous mutation conferring resistance to streptomycin. *FRT* refers to the FLP site-specific recombination site.

[b]Tn*10* transposon is located in the intergenic region *ybdB-ybfA* on the F plasmid.

[c]*sfgfp* gene encodes the superfolder Green Fluorescent Protein sfGFP

**Table S1. Strains list**

| Name | Construct and Usage | Source or reference |
|---|---|---|
| pCP20 | Flp expression plasmid | Datsenko *et al.,* 2000 |
| pKD3 | Carries *FRT-Cm-FRT* used for λred integration ; CmR | Datsenko *et al.,* 2000 |
| pKD4 | Carries *FRT-kan-FRT* used for RecET integration; KnR | Datsenko *et al.,* 2000 |
| pmCherry-*ParB* (pSN70) | IPTG inducible expression of N-terminal fusion mCherry-$ParB_{PMT1}$ | Nolivos *et al.,* 2019 |
| pR6K-sfGFP | Carries *sfgfp-FRT-kan-FRT* used for several C-terminal fusion by λred | Nolivos *et al.,* 2019 |
| pR6K-$P_{biofab}$-sfGFP | Carries *sfgfp* gene under the *Biofab* promotor | Reuter *et al.,* 2020 |
| pR6K- $P_{biofab}$-mCherry | Carries *mcherry* gene under the *Biofab* promotor | Reuter *et al.,* 2020 |
| pR6K-$P_{tac}$-mCherry | Carries *mcherry* gene under the *tac* promotor | Replacement of *pBiofab* in pR6K-pBiofab-mCherry by *ptac* |
| pUA66-$P_{sulA}$-GFP-kan | SOS inductible expression of *gfp-frt-kan-frt* fusion *sulA* promotor with *pSC101* oriV | Gift from L. Van Melderen |
| pUA66-$P_{sulA}$-GFP-gm | SOS inductible expression of *gfp-frt-gm-frt* fusion *sulA* promotor with *pSC101* oriV | Replacement of *frt-kan-frt by frt-gm-frt* |
| pNF02-$P_{pspA}$-mNG | Psp inducible expression of *mneongreen* fusion *pspA* promoter on mini-F plasmid | Rousseau *et al.,* 2023 |
| pNF02-$P_{micA}$-mNG | SigmaE inducible expression of *mneongreen* fusion *micA* promoter on mini-F plasmid | Rousseau *et al.,* 2023 |
| pNF02-$P_{cpxP}$-mNG | Cpx inducible expression of *mneongreen* fusion *cpxP* promoter on mini-F plasmid | Rousseau *et al.,* 2023 |
| pNF02-$P_{mdtA}$-mNG | Bae inducible expression of *mneongreen* fusion *mdtA* promoter on mini-F plasmid | Rousseau *et al.,* 2023 |
| pNF02-$P_{rcsA}$-mNG | Rcs inducible expression of *mneongreen* fusion *rcsA* promoter on mini-F plasmid | Rousseau *et al.,* 2023 |
| pAC2 | Carries *traS* gene under the *Trc* promotor | Insertion of *traS* in pTrc99a |

| | | |
|---|---|---|
| pAC3 | Carries *traT* gene under the *Trc* promotor | Insertion of *traT* in pTrc99a |
| pAC4 | Carries *traS* and *traT* gene under the *Trc* promotor | Insertion of *traST* in pTrc99a |
| pAC13 | Psp inducible expression of *mneongreen* fusion *pspA* promoter with *pSC101* oriV | Replacement of $P_{sulA}$-*sfgfp* from pUA66-$P_{sulA}$-sfGFP-gm with $P_{pspA}$-mneongreen using Gibson assembly |
| pAC14 | SigmaE inducible expression of *mneongreen* fusion *micA* promoter with *pSC101* oriV | Replacement of $P_{sulA}$-*sfgfp* from pUA66-$P_{sulA}$-sfGFP-gm with $P_{micA}$-mneongreen using Gibson assembly |
| pAC15 | Cpx inducible expression of *mneongreen* fusion *cpxP* promoter with *pSC101* oriV | Replacement of $P_{sulA}$-*sfgfp* from pUA66-$P_{sulA}$-sfGFP-gm with $P_{cpxP}$-mneongreen using Gibson assembly |
| pAC16 | Bae inducible expression of *mneongreen* fusion *mdtA* promoter with *pSC101* oriV | Replacement of $P_{sulA}$-*sfgfp* from pUA66-$P_{sulA}$-sfGFP-gm with $P_{mdtA}$-mneongreen using Gibson assembly |
| pAC17 | Rcs inducible expression of *mneongreen* fusion *rcsA* promoter with *pSC101* oriV | Replacement of $P_{sulA}$-*sfgfp* from pUA66-$P_{sulA}$-sfGFP-gm with $P_{rcsA}$-mneongreen using Gibson assembly |
| pmob | pSEVA231-*oriT-F* | Reuter *et al.,* 2020 |
| **F-Tn*10* conjugative plasmid (from K603) derivatives** | | |
| F*wt* | F-Tn*10* with *parSPMT1* inserted at the intergenic *ygeB-ygfA locus* | Fig.1-6, Fig. S1-S4 |
| F*ΔtraS* | F-Tn*10 parSPMT1* with *traS* deletion | Fig.1, 2, 4, Fig. S1 |
| F*ΔtraT* | F-Tn*10 parSPMT1* with *traT* deletion | Fig.1, 2, 4, Fig. S1 |
| F*ΔtraST* | F-Tn*10 parSPMT1* with *traS-traT* deletions | Fig.1-6, Fig. S1-S4 |
| F*ΔtraIST* | F-Tn*10 parSPMT1* with *traI* and *traS-traT* deletions | Fig.1-3, Fig. S1, S3-S4 |
| F*ΔtraAST* | F-Tn*10 parSPMT1* with *traA* and *traS-traT* deletions | Fig.1-2, Fig. S1, S3-S4 |

| | | |
|---|---|---|
| **FΔ*traST*$^{GFP+}$** | F-Tn*10 parSPMT1-ΔtraST* with insertion $P_{biofab}$-*sfgfp* in the *repE-sopA* intergenic region | Fig. 4 |
| **FΔ*traST*$^{RFP+}$** | F-Tn*10 parSPMT1-ΔtraST* with insertion $P_{tac}$-*mcherry* in the *repE-sopA* intergenic region | Fig. 4, 5 |
| **Fwt$^{GFP+}$** | F-Tn*10 parSPMT1* with insertion $P_{biofab}$-*sfgfp* in the *repE-sopA* intergenic region | Fig.4, 5 |
| **F*wt*$^{RFP+}$** | F-Tn*10 parSPMT1* with insertion $P_{tac}$-*mcherry* in the *repE-sopA* intergenic region | Fig.4, 5 |
| **FΔ*oriT*** | F-Tn*10 parSPMT1* with *oriT* deletion | Fig.5, Fig.S4 |
| **FΔ*oriT*Δ*traST*** | F-Tn*10 parSPMT1* with *oriT* and *traST* deletion | Fig.5, Fig.S4 |

[a] *sfgfp* gene encodes the superfolder Green Fluorescent Protein sfGFP

Table S2. Plasmids used in this study

| Name | Sequence | Construct |
|---|---|---|
| OL117 | GAGCATAGCGAGCGAACTGGCGAGGAAGCAAAGAAGAACTCATATGAATATCCTCCTTAG | λred *pBiofab-sfgfp-FRT-kan-FRT* construct at F plasmid intergenic *repE locus*. PCR on pR6K-pBiofab-sfGFP plasmid. |
| OL383 | TCTGGAAGATTTTGCCGAACCACAAATGACGTTGTCGCGCCATATGAATATCCTCCTTAG | λred *ptac-mcherry-FRT-kan-FRT* construct at the endogenous *ilvA locus*. PCR on pR6K-ptac-mcherry plasmid. |
| OL688 | TTTCTTCTTGCGCTGAGCGTAAGAGCTATCTGACAGAACCCGACTGGAAGCATCGATAG | λred *pBiofab-sfgfp-FRT-kan-FRT* construct at F plasmid intergenic *repE locus*. PCR on pBiofab-pR6K-sfGFP plasmid. |
| OL733 | GTTATCAAGAGTAAAATAAAAGATATTAGAGAGTAAATATCATATGAATATCCTCCTTAG | λred Δ*traT::cat* construct at the endogenous F plasmid *locus*. PCR on pKD3 |
| OL734 | GTCAGTCAGGAGGCCGGTCAGACCAGCCTCCGGAAGATAAGTGTAGGCTGGAGCTGCTTC | |
| OL737 | AAGAATAATCATAACATGTTGGGTAGGGTATGGAGAGACCCATATGAATATCCTCCTTAG | λred Δ*traS::cat* construct at the endogenous F plasmid *locus*. PCR on pKD3 |
| OL738 | GTTTTTTTGTTTTCATCATATATTTACTCTCTAATATCTTGTGTAGGCTGGAGCTGCTTC | |
| OL743 | gaaaaatgcctgatagcgcttcgcttatcaggcctacccgGACAGGTTTCCCGACTGGAA | λred *ptac-mcherry-FRT-kan-FRT* construct at the endogenous *ilvA* |

| | | |
|---|---|---|
| | | *locus*. PCR on pR6K-ptac-mcherry plasmid. |
| OL767 | CACTCTAGTTTTATTCATTTATCCGAAATTGAGGTAACTT CATATGAATATCCTCCTTAG | λred Δ*traA::kan* construct at the endogenous F plasmid *locus*. PCR on pKD4 |
| OL768 | TTAAGTTTATTCTCGTCTCCCGACATCGTTTTATTTCCTG GTGTAGGCTGGAGCTGCTTC | |
| OL817 | atttcttcttgcgctgagcgtaagagctatctgacagaacGACAGGTT TCCCGACTGGAA | λred *pBiofab-sfgfp-FRT-kan-FRT* construct at F plasmid intergenic *repE locus*. PCR on pR6K-pBiofab-sfGFP plasmid. |
| OL944 | GTGGGATTGATGCCGGGATATGTCAAAGGGATATACGT TTCATATGAA TATCCTCCTTAG | λred Δ*traI::kan* construct at the endogenous F plasmid *locus*. PCR on pKD3 |
| OL945 | GTTCGTGTTATCCGTTGTCATCAGCGTTTGTCTTCCTGTA GTGTAGGC TGGAGCTGCTTC | |

Table S3. PCR primers used for strain and plasmid constructions
